# Gut bacteria impact host uric acid burden and its association with atherosclerosis

**DOI:** 10.1101/2022.12.12.520169

**Authors:** Kazuyuki Kasahara, Robert L. Kerby, Qijun Zhang, Meenakshi Pradhan, Margarete Mehrabian, Aldons Lusis, Göran Bergström, Fredrik Bäckhed, Federico E. Rey

**Affiliations:** Department of Bacteriology, University of Wisconsin-Madison, Madison, Wisconsin, USA; Lee Kong Chian School of Medicine, Nanyang Technological University, Singapore, Singapore; Department of Molecular and Clinical Medicine, Wallenberg Laboratory, Institute of Medicine, University of Gothenburg, Gothenburg, Sweden; Division of Cardiology, Department of Medicine, David Geffen School of Medicine, University of California-Los Angeles, Los Angeles, USA; Region Västra Götaland, Sahlgrenska University Hospital, Department of Clinical Physiology, Gothenburg, Sweden; Novo Nordisk Foundation Center for Basic Metabolic Research, Faculty of Health Sciences, University of Copenhagen, Denmark

## Abstract

Humans with metabolic and inflammatory diseases, including atherosclerosis harbor dysbiotic gut communities. However, the microbes and microbial pathways that influence disease progression remain largely undefined. Here, we show that variation in atherosclerosis burden is in part driven by the gut microbiota and it is associated with circulating levels of the proinflammatory molecule uric acid both in mice and humans. We identify bacterial taxa present in the gut spanning multiple phyla, including *Bacillota* (Firmicutes), *Fusobacteriota* and *Pseudomonadota* (Proteobacteria), that use uric acid and adenine– a key precursor of nucleic acids in intestinal cells, as carbon and energy sources anaerobically, and uncover a gene cluster encoding key steps of purine degradation that is widely distributed among gut dwelling bacteria. Furthermore, we demonstrate that colonization of germ-free mice with purine-degrading bacteria modulates levels of uric acid and other purines in the gut and systemically. Altogether this work demonstrates that gut microbes are important drivers of host global purine homeostasis and uric acid levels, and suggests that gut bacterial catabolism of purines may represent a novel mechanism by which the gut microbiome influences host health.

## Introduction

Metabolic disorders including obesity, type 2 diabetes, and atherosclerosis have historically been viewed as lipid conditions primarily driven by overindulgence of calorie dense foods. However, it is now widely appreciated that chronic inflammation plays a central role in the development and progression of these disorders^1^. Atherosclerosis, the leading cause of cardiovascular disease (CVD), is characterized by vascular inflammation, and it is influenced by multiple genetic and environmental factors^1–3^. Large-scale genome-wide analyses in human populations have identified over 100 loci significantly associated with atherosclerosis itself^4^ and hundreds of loci for traits associated with atherosclerosis such as plasma lipids, obesity, and diabetes^5–7^. Nevertheless, while atherosclerosis has a significant genetic component, the environment, especially diet also plays a major role in its progression. Furthermore, several recent studies have provided evidence suggesting that dietary contributions to disease progression are often mediated by the gut microbiome^8–10^.

The gut microbiome exerts profound influence on metabolism and inflammatory diseases^11,12^. Diet and host-derived factors modulate the composition of the gut microbiome, which in turn transforms dietary components consumed by the host, generating bioactive molecules that interact with the immune system and virtually every organ in the host’s body, including the vascular system. Changes over the last century in food production, dietary habits, antibiotic usage, and lifestyle have caused major changes in the microbiome and have greatly impacted human health in discordant ways^13–15^. During the past decades the prevalence of acute infectious diseases has decreased, while it increased for chronic inflammatory diseases. Furthermore, several diet-derived gut bacterially-produced metabolites have been uncovered as potential drivers of metabolic and cardiovascular ailments. These metabolites constitute a direct link between environmental exposures and host cellular function and encompass several uremic toxins (i.e., waste products that cannot be eliminated properly by subjects with impaired kidney function), including *p*-cresyl sulfate, indoxyl sulfate, trimethylamine-N-oxide (TMAO), and phenylacetylglutamine^16^. For example, dietary choline, betaine and carnitine serve as substrates for TMAO production, which is generated in the liver from gut bacterially-produced trimethylamine (TMA)^8^. TMAO enhances inflammation and aortic thrombosis in mice and it is associated with CVD risks in humans^8,17,18^. More recently it was found that phenylacetylglutamine, synthesized by the microbiota from dietary protein, enhances platelet activation and thrombosis via host G-protein coupled receptors^19^. Together, this evidence supports the notion that gut bacteria metabolism contributes to CVD-related traits by modulating abundance of uremic toxins in circulation.

Uremic toxins impair several biological functions, contributing to several symptoms observed in subjects with chronic kidney disease^16,20^. Several purines-derived metabolites, including xanthine, hypoxanthine and uric acid (UA) are also considered uremic toxins^21^. UA–the end product of metabolic breakdown of purines in humans, is mostly studied for the complications it causes when its concentration reaches saturation levels, forming pro-inflammatory crystals that deposit in joints (e.g. gout). However, a recent study showed that concentrations of UA within the solubility range can promote atherosclerosis via induction of AMP-activated protein kinase (AMPK)-mediated inflammation^22^. UA exacerbates inflammation, endothelial dysfunction, increases the renin-angiotensin-aldosterone system activity^21^ and it is increased in patients with hypertension and heart failure^20^. Moreover, several studies have suggested that pharmacological interventions effective at reducing UA production or increasing its excretion in hyperuricemic patients improves cardio-renal outcomes^23,24^, whereas others have not found such benefits^25^. While the kidneys play a major role in regulating levels of UA in circulation, a significant fraction of this metabolite is secreted into the intestine^26^ and a recent metagenomic study identified bacterial pathways associated with blood levels of UA^27^. However, no causal relationships have been established between abundance of this uremic toxin and specific gut bacteria.

We sought to examine the role of the gut microbiome on atherosclerosis and identify potential microbial pathways that contribute to disease burden. First, we transplanted microbial communities derived from mouse strains with disparate atherosclerosis phenotypes into germ-free (GF) *Apolipoprotein E* knockout (*ApoE* KO) mice. We found that microbial-driven variation on atherosclerosis progression was associated with abundance of purine metabolites including UA. We also observed that this pro-inflammatory metabolite was associated with atherosclerosis burden and gut microbial features in a human cohort. We identified taxa able to degrade purines anaerobically, uncovered a gene cluster encoding key components needed for anaerobic purine degradation, demonstrated factors affecting its activity, and showed that colonization with taxa containing this locus lowered purines in the gut and UA systemically in mice. Altogether this work strengthens the connection between gut microbes and atherosclerosis and provides new insights into how bacterial metabolism influences host biology.

## Results

### Gut microbes modulate atherosclerosis progression and plasma metabolites associated with disease burden

Previous work revealed a large degree of variation in atherosclerosis burden among 100 inbred strains of mice from the Hybrid Mouse Diversity Panel (HMDP), which harbor distinct microbial communities^28,29^. We hypothesized that the gut microbiome contributed to the variation observed in disease progression among HMDP strains. We transplanted cecal samples from four HMDP strains into GF *ApoE* KO recipient mice. We selected two strains that exhibited large atherosclerotic lesions (AXB10/PgnJ and BXD5/TyJ) and two strains that showed little signs of disease (BTBR T+tf/J and BXA8/PgnJ), hereinafter referred as “AXB10”, “BXD5”, “BTBR”, and “BXA8”, respectively. Transplanted mice were maintained on a chow diet supplemented with 0.2% cholesterol for 8 weeks. After this period, atherosclerotic lesions, gut microbiome composition, and disease biomarkers were evaluated (**Fig 1A, Suppl. Fig 1**). We found that mice colonized with cecal communities from HMDP donors prone to atherosclerosis development (i.e., AXB10 and BXD5) exhibited larger lesions compared to recipient mice colonized with samples from donors that showed little signs of atherosclerosis (i.e., BTBR and BXA8). These results support the notion that the gut microbiome contributes to the development of atherosclerosis and possibly to the variation in disease burden observed among the HMDP strains (**Fig 1B-G**). Neither traditional CVD risk factors such as body weight and cholesterol, nor previously identified gut microbiota-derived metabolites including lipopolysaccharides (LPS), TMAO, and short-chain fatty acids explained the differences in atherosclerosis burden observed among the transplanted mice (**Suppl. Fig 1**).

**Figure 1.**
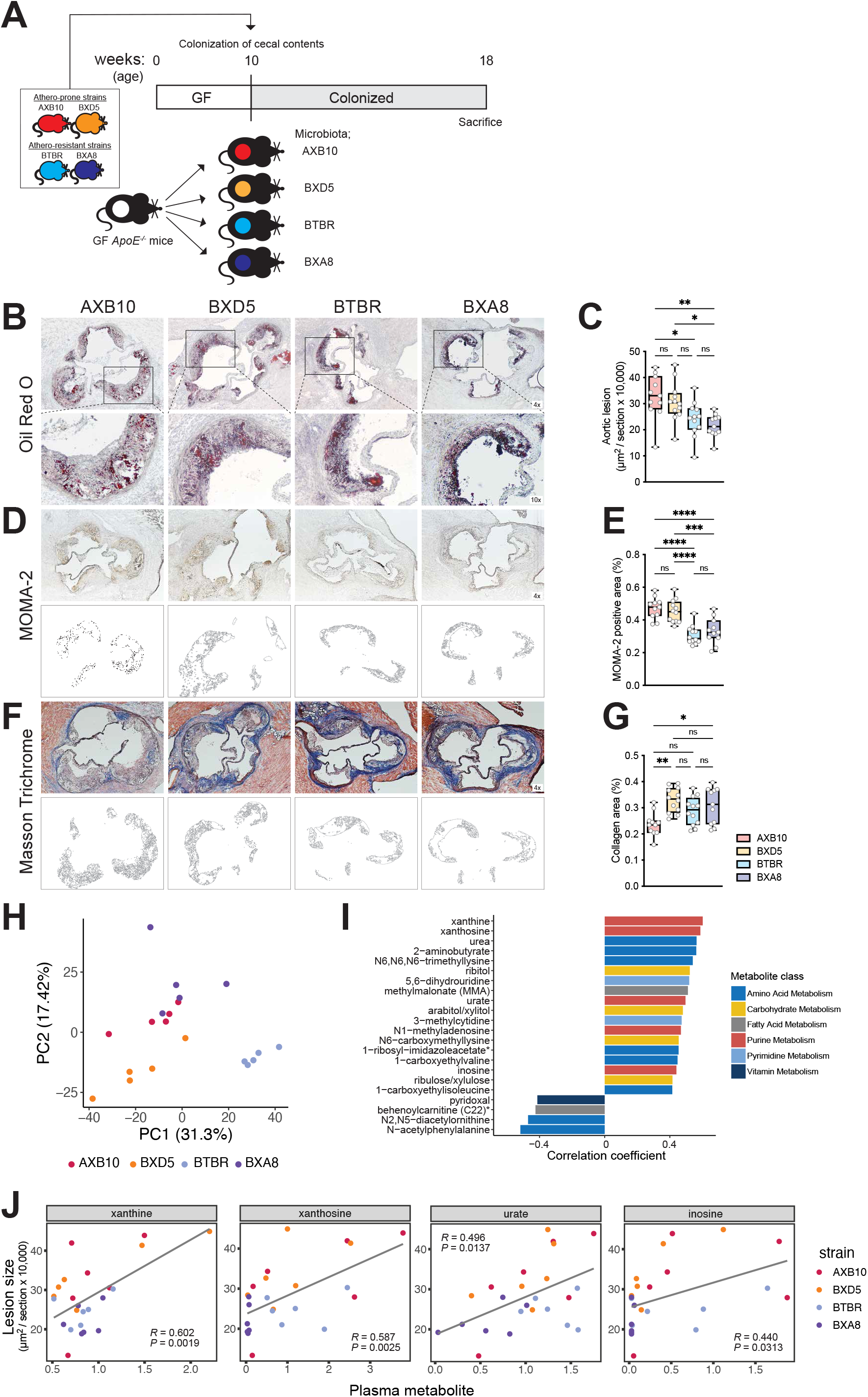
Plasma levels of purines are associated with atherosclerosis burden in transplanted gnotobiotic *ApoE* KO mice. **A)** Experimental design. **B-G)** Representative sections and quantitative analysis of Oil Red O staining (B, C), MOMA-2 staining (D, E), and Masson’s trichrome staining (F, G) in the aortic sinus (n=11 for AXB10, n=12 for BXD5, n=12 for BTBR group, n=11 for BXA8). The data are expressed as box-and-whisker plots with individual data points, where the boxes indicate the median values and the interquartile ranges and the whiskers represent the minimum and maximum values. Significance was calculated by one-way ANOVA with the Tukey post-tests is indicated as follows: *, *p*-value <0.05; **, *p*-value of <0.01; ****, *p*-value of <0.0001. **H)** Principal component analysis of gut microbial functions from transplanted mice as determined by metagenomic analysis. **I)** Plasma metabolites positively or negatively associated with atherosclerotic lesion size, according to Spearman correlation analysis. **J)** Scatterplots showing associations between purines (relative mass spectrometry scaled intensities) and atherosclerosis lesion size (x10^4^ μm^2^). GF; germ-free, *ApoE*; *Apolipoprotein E*, Chol; cholesterol, MOMA; monocytes and macrophages.

Shotgun metagenomic analyses of cecal contents from the transplanted mice identified 1649 functional features and 52 bacterial taxonomic features. Principal component analysis of functional features show distinct clustering by donor strain (*p*-value <0.001, PERMANOVA) suggesting unique gut bacterial function profiles for each of the four HMDP strains used (**Fig. 1H**). Pearson correlation analysis identified several bacterial functions associated with atherosclerosis lesion size among recipient mice (**Suppl. Fig. 2A, B**). These included several functions related to production and conversion of purines. We also observed that bacterial pathways including energy metabolism, amino acid metabolism were enriched among functions positively correlated with atherosclerosis lesion size, although these did not survive multiple hypothesis correction.

To further investigate whether microbiota transplants impacted circulating metabolites associated with disease, we performed metabolome analysis of plasma samples using Ultra-high Performance Liquid Chromatography (UPLC) - Tandem Mass Spectrometer (MS/MS). A total of 682 metabolites were measured (**Suppl. Table 1**). Pearson correlation analyses identified purine metabolites including xanthine, xanthosine, inosine, and UA, positively associated with atherosclerotic lesions size (**Fig. 1I, J**). Altogether, these results suggested that gut microbes influence atherosclerosis progression and abundance of purines in blood of transplanted mice.

### Serum UA is correlated with gut microbial features and subclinical atherosclerosis in a human cohort

UA is the end product of purine metabolism in humans, and it has been shown to cause inflammation^22,30,31^, induce endothelial dysfunction^32^ and stimulate smooth muscle cell proliferation^33^. We explored associations between atherosclerosis, gut bacteria, and UA in a human cohort previously characterized for gut microbiome and glucose homeostasis (n=998)^34^. Coronary artery calcium (CAC) score measurements were assessed for disease burden. Calcification of arteries is an accepted proxy for estimating overall plaque burden of atherosclerosis^35^. We first classified individuals based on their CAC score status: CAC score = 0 (i.e., no detectable vascular calcification, n=492) *vs.* CAC >0 (n=497). Logistic regression analysis revealed that distribution of UA concentrations was significantly different between individuals from these two groups (**Fig. 2A**) with individuals with CAC > 0 showing higher mean and median UA levels. Furthermore, Spearman correlation analysis for individuals with CAC score >0 showed a significant positive association between UA and CAC score (rho coefficient = 0.14, *p*-value <0.001, **Fig. 2B**). We then applied extreme gradient boosting (XGBoost) regression to identify gut bacterial taxa correlated with UA levels. XGBoost is a decision-tree-based ensemble machine learning algorithm that uses the gradient boosting method. The top 10 features associated with UA levels are shown in **Suppl. Table 2** after adjusted for covariates (CACS, BMI, gender, triglycerides and HbA1c) in a mixed linear regression model. Interestingly, we found multiple taxa within the *Clostridia* class that were negatively associated with levels of UA. Altogether these results suggest that the gut microbiome, particularly taxa within the *Clostridia*, may influence UA levels. This data is also consistent with gnotobiotic mouse work reported above and previous work connecting UA with CVD in humans.

**Figure 2.**
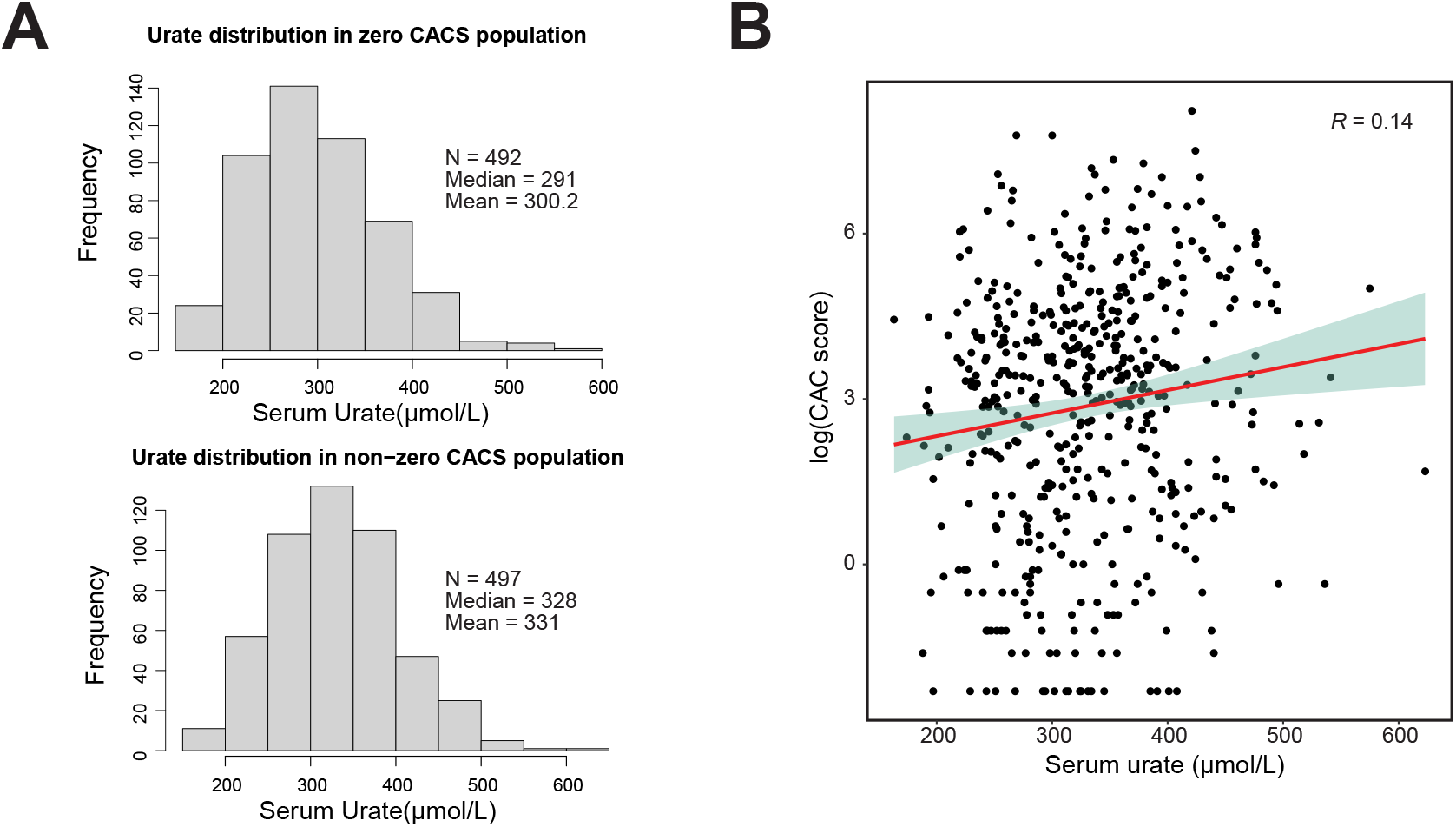
Plasma uric acid levels are positively associated with Coronary Artery Calcium (CAC) score in a human cohort. **A)** Distribution of uric acid levels in serum from individuals with CAC score=0 and CAC score >0. **B)** Spearman correlation analysis between serum uric acid levels and CAC score.

### Gut microbiome modulates purines in cecum and circulation

We next investigated whether the gut microbiome modulated abundance of purines in the intestine and circulation. We quantified purine related metabolites, including nucleotides, nucleosides and nucleobases by LC-MS/MS in cecal contents and plasma from GF mice and conventionally-raised (Conv) animals (**Suppl. Table 3, 4**). We found that most purines were decreased in the cecal contents from GF mice compared to Conv mice, with a few exceptions being increased in GF mice, such as UA and allantoin– both waste products of purine metabolism in mice (**Fig. 3A, B**).

**Figure 3.**
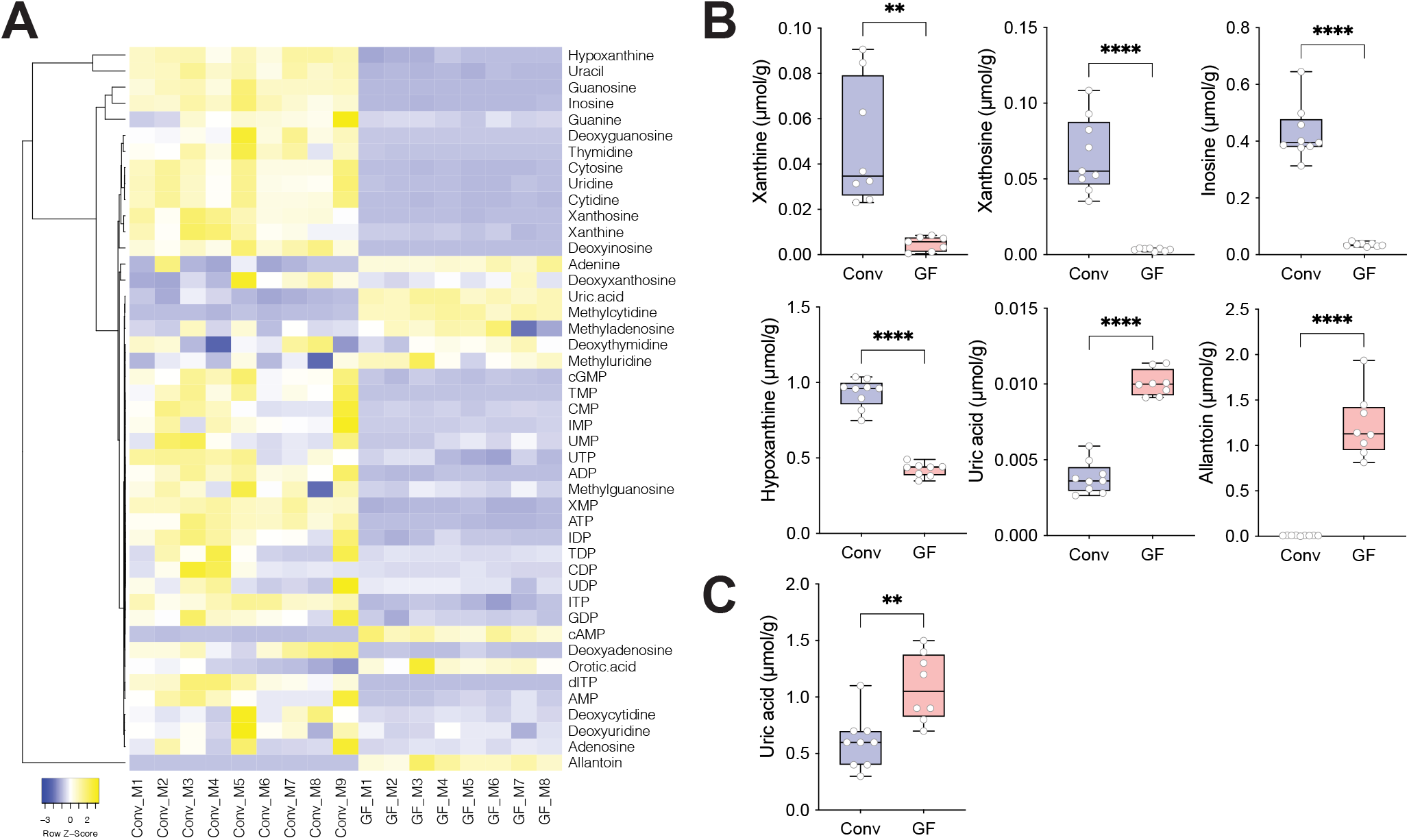
Gut microbiome modulates purines in cecum and circulation. **A)** Heatmap of purine metabolites in cecal contents from Conv (n=9) and GF (n=8) mice. **B)** Values for xanthine, xanthosine, inosine, hypoxanthine, uric acid, and allantoin in cecal contents measured by LC/MS/MS and **C**) and plasma uric acid levels analyzed by enzymatic assay are shown using box-and-whisker plots with individual data points, where the boxes indicate the median values and the interquartile ranges and the whiskers represent the minimum and maximum values. Significance was calculated by unpaired two-tailed Student’s t-test and is designated as follows: **, *p*-value of <0.01; ****, *p*-value of <0.0001. Conv; Conventionally raised, GF; germ-free.

Partial Least Squares-Discriminant Analysis (PLS-DA) analysis of cecal purines showed separation of the two groups by principal components 1 and 2. While the separation for plasma samples was less evident (**Suppl. Fig. 3A, B**), we found that GF mice had significantly increased UA levels in plasma compared to Conv mice (**Suppl. Fig. 3A, C**). This result was confirmed using an enzymatic assay to quantify UA (**Fig. 3C**). Altogether these results suggested that the gut microbiome modulates abundance of purines both in the gut and systemically.

### Microbes degrade UA and adenine anaerobically

The intestine is a key organ for purine homeostasis. Dietary purines are absorbed in the gut, resident microbes produce and recycle purines needed for their anabolism and ~30% of the UA generated by the body is secreted into the intestine^36,37^. We hypothesized that bacteria influence UA, by consuming different purines from diet and UA. While this notion has been previously discussed, isolates from the human or mouse gut able to degrade these metabolites anaerobically have not been identified. We attempted anaerobic enrichments using fecal slurries (human samples) on media supplemented with UA as the primary source of carbon and energy. Culture media included non-fermentable acetate (often beneficial for butyrate-producing Firmicutes) plus 0.1% yeast extract as the sole complex nutrient. Enrichments were plated and colonies isolated on bilayer agar plates bearing a top agar layer supplied with saturating amounts of UA or other purines as described in the Materials and Methods. We isolated several Firmicutes and Proteobacteria including species identified as *Enterocloster bolteae* and *Escherichia coli* by 16S rRNA gene sequencing. Using the same medium we verified growth and substrate utilization by *E. bolteae* ATCC BAA-613—the species type strain, which has been previously sequenced. We observed that cultures of this strain supplemented with 12 mg UA degraded 49.6 μmoles substrate in a 24-hour period, with accumulation of 106.9 μmoles acetate, a known fermentation product of this organism. As analyzed by High Pressure Liquid Chromatography (HPLC) and headspace Gas Chromatography (GC), significant ethanol, formate, propionate, lactate and butyrate accumulation were not observed under this condition.

We then expanded our search for bacteria with this capability and by using methods described above screened a culture collection of 34 isolates encompassing gut-dwelling bacteria from six phyla (**Suppl. Fig. 4;** a representative subset of these species is shown in **Fig. 4).** The screen utilized monolayer plates containing no nitrogen (other than 0.1% yeast extract) or fermentable carbon or energy source, the same medium supplemented with soluble substrates (glucose or allantoin), or bilayer plates (described above, see also Materials and Methods) containing insoluble purines (UA or adenine). These assays demonstrated the anaerobic allantoin-, UA- and adenine-dependent growth characteristics of these strains as evident by growth of the applied bacterial patch and a zone of disappearance of the insoluble UA or adenine substrates (**Fig. 4, Suppl. Fig. 4**). Our screen showed evidence of UA and adenine utilization among the *Bacillota* (Firmicutes), *Fusobacteriota* and *Pseudomonadota* (Proteobacteria) phyla (**Fig. 4**, **Suppl. Fig. 4**), although this property was not universal among strains belonging to these phyla. Of particular note, these assays showed different purine-utilization capacities among UA degrading strains: (i) *E. coli* MS 200-1 and *E. coli* I-11 showed more efficient UA utilization compared to the typically analyzed *E. coli* K12 strain (**Fig. 4**); (ii) allantoin supported growth of *E. bolteae* and *E. coli* but not medically-relevant *Clostridiodes difficile* or *Edwarsiella tarda* (**Fig. 4**); (iii) adenine supported the growth of several strains of Proteobacteria (*E. coli* MS200-1, *E. coli* I-11, *E. tarda*) and Firmicutes (*E. bolteae, E. asparagifome*), but not *C. difficile* or *C. sporogenes* (**Fig. 4**, **Suppl. Fig. 4**) (iv) consistent with previous work, growth of Proteobacteria strains on UA was enhanced by the addition of formate^38^ (**Suppl. Fig. 4**), while the addition of formate had more modest effects, if any, for strains from other phyla tested. It is also important to note that none of the six *Bacteroides* strains tested showed any growth on UA, allantoin or adenine (**Fig. 4**, **Suppl. Fig. 4**).

**Figure 4.**
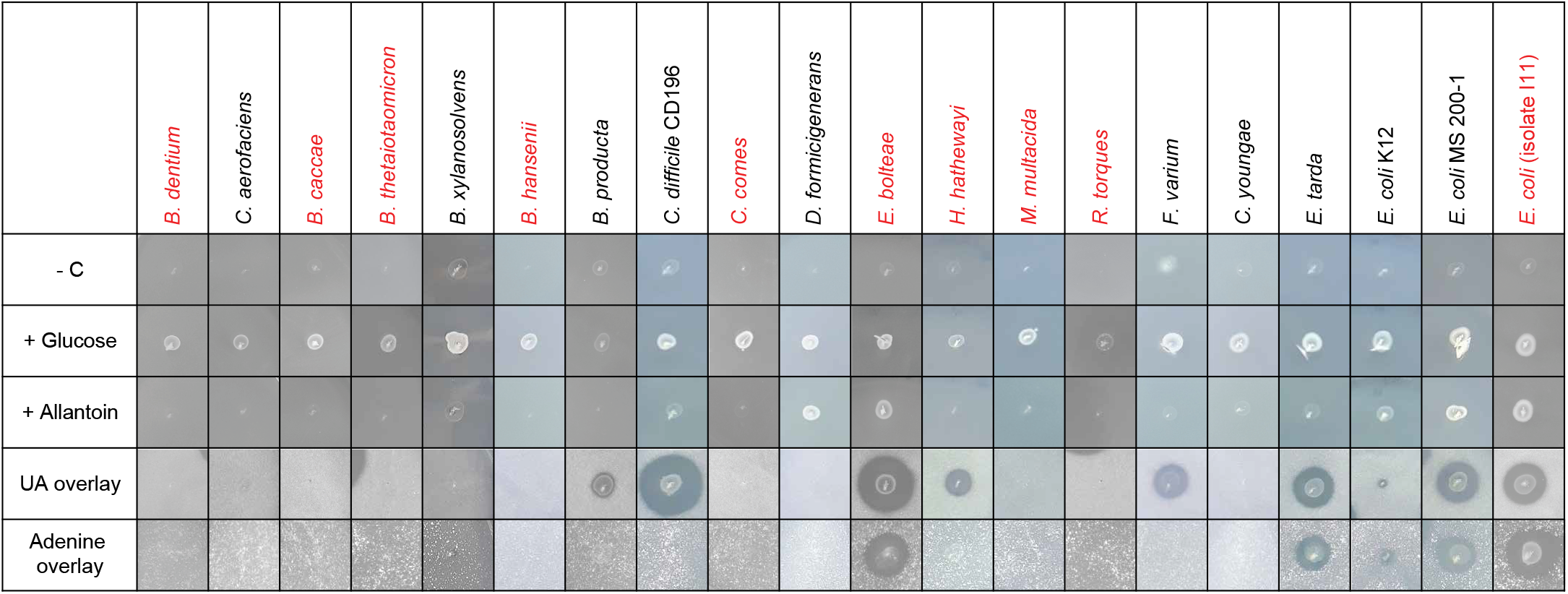
Gut bacterial isolates use purines as carbon and energy source. Anaerobic growth of bacterial strains on plates containing soluble (glucose, allantoin), and insoluble (uric acid and adenine) substrates. As detailed in the Materials and Methods, plates were inoculated with 4 μl of dense overnight cultures grown in rich medium then incubated for 2 days [no substrate (-C), glucose/NH4, allantoin or uric acid conditions] or 7 days (adenine). Growth is indicated by the appearance of cell patches and a zone of clearing for the overlay plates. Tested species included *Collinsella aerofaciens*, *Bacteroides caccae*, *Bacteroides dorei*, *Bacteroides finegoldii*, *Bifidobacterium dentium*, *Blautia hansenii*, *Blautia producta*, *Clostridium sporogenes*, *Clostridium symbiosum*, *Coprococcus comes*, *Dorea formicigenerans*, *Enterocloster asparagiforme*, *Enterocloster bolteae*, *Erysipelatoclostridium ramosum*, *Hungatella hathewayi*, *Fusobacterium varium*, *Citrobacter youngae* and *Escherichia coli* MS 200-1. Details about strains are specified in Supplemental Table 10, and a summary of all tested strains is presented in Supplemental Figure 4. Strains indicated in red were used for colonization of gnotobiotic mice.

In multiple organisms the ability to use UA was diminished or eliminated in media containing glucose or fructose, consistent with regulation by catabolite repression (**Suppl. Fig. 5**). Another factor affecting UA utilization—at least for Proteobacteria species—was the requirement for Molybdenum (Mo) and Selenium (Se) supplementation. Although no attempt was made to limit trace minerals—i.e., the medium contained a high amount of phosphate buffer plus low levels of yeast extract and cysteine (a possible source of Se contamination)^39^ and the inocula were prepared in rich medium—a demonstrable requirement for micromolar additions of Mo and Se was evident for growth on UA by both species of Proteobacteria tested (**Suppl. Fig. 5**). However, these effects were not observed with the two Firmicutes tested, *C. difficile* and *E. bolteae* (**Suppl. Fig. 5**). Additional experiments did not show an effect of Co, Mn, Ni, W, or Zn supplementation on UA utilization by any of the tested strains (*E. bolteae, C. difficile, E. coli* MS 200-1, *E. tarda*). Altogether, the results presented above suggested that common gut bacteria can use purines for carbon and energy, and that availability of other carbon sources and metals modulate this process.

### Purine-degrading bacteria modulate abundance of purines in cecum and circulation

To test the *in vivo* functions of purine-degrading bacteria (PDB) identified above, we created synthetic bacterial communities that varied in their capacity to degrade UA and used them to colonize GF mice. GF mice were colonized with a core community which included seven species that do not degrade purines *in vitro*, *Bacteroides caccae*, *Bifidobacterium dentium*, *Blautia hansenii*, *Bacteroides thetaiotaomicron*, *Coprococcus comes*, *Mitsuokella multacida*, and *Ruminococcus torques* or the core community plus three PDB, including *E. bolteae*, *Hungatella hathewayi,* and *E. coli* isolate l-11 (**Fig. 5A**). *In vitro* tests of UA utilization for each of the strains used in this community is shown in **Fig. 4** (highlighted in red), and their combined activities are indicated in **Fig. 5A**. A third group of animals remained germ-free throughout the experiment. The engrafted bacterial communities were analyzed by COPRO-Seq (*co*mmunity *pro*filing by *seq*uencing) analysis^40^. All taxa included in the communities successfully colonized the gut of GF mice. *E. bolteae* was the most abundant among the three PDB (relative abundance: *E. bolteae* 19.5%, *H. hathewayi* 5.6% and *E. coli* 0.9%, **Fig. 5B**). Assays of *in vitro* UA utilization by fecal samples collected from colonized mice showed that colonization with PDB was required for UA degradation. We next performed targeted purine quantification in cecal contents and plasma (**Suppl. Table 5, 6**). Global analysis of the data showed distinct patterns for cecal purine-related metabolites between GF mice and mice with the “core” or the “core plus PDB”, where nucleosides were increased and nucleotides were decreased in the cecum of GF mice (**Fig. 5C**). Surprisingly, mice colonized with the “core plus PDB’’ community showed significantly higher levels of UA in the cecum compared to mice colonized with the “core” community, while cecal levels of other purines/nucleosides including hypoxanthine, adenosine and allantoin were significantly reduced in mice co-colonized with the three PDB (**Fig. 5D**). It is important to note that these metabolites were detected at significantly higher concentrations in the gut of mice relative to UA (**Suppl. Table 5)**. PLS-DA analysis separated plasma samples from the core plus PDB to the ones from the other two groups (**Suppl. Fig. 6A, B**). Colonization with core plus PDB resulted in significantly lower levels of several purine metabolites in plasma, including UA (**Suppl. Fig. 6A, C**). Plasma UA results were also confirmed by an enzymatic assay (**Fig. 5E**). Altogether these results suggested that purine degrading bacteria impacted levels of several purines in the gut and specifically UA both locally and in circulation.

**Figure 5.**
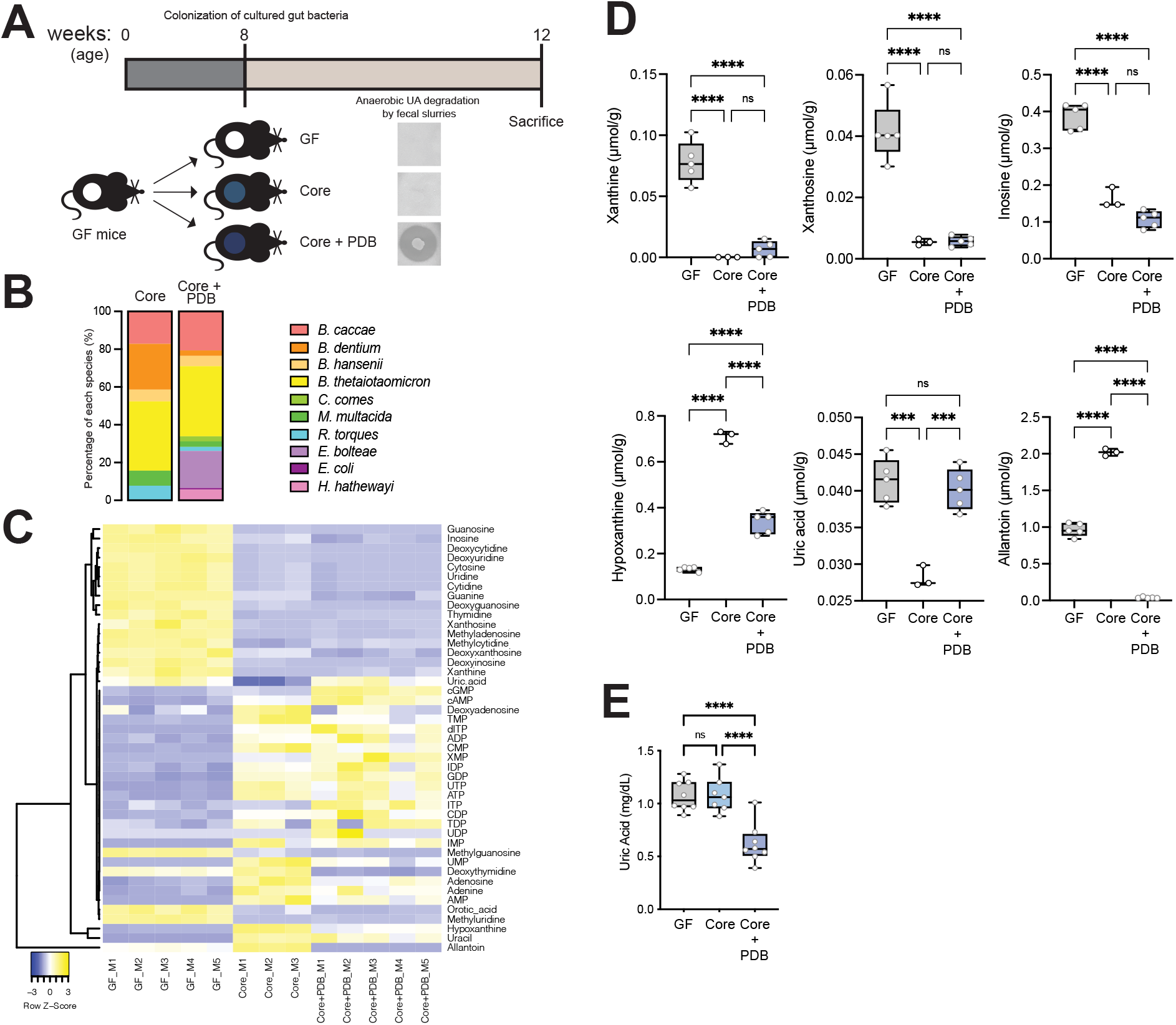
Purine-degrading bacteria modulate abundance of purines in cecum and circulation. **A)** Experimental design. Anaerobic uric acid degradation by fecal samples from different groups is indicated using uric acid overlay plates as detailed in Material and Methods. **B)** Community profiling by sequencing (COPRO-Seq) analysis of fecal samples from gnotobiotic B6 mice colonized with the ‘core’ community (n=4) or the ‘core plus purined degrading bacteria’ community (n=3). The bar charts show the abundance of each species in each community. **C)** Heatmap of purine metabolites in cecal contents from GF (n=5), ‘core’ (n=3) and ‘core plus PDB’ (n=5) mice analyzed by LC-MS/MS. **D)** Values for xanthine, xanthosine, inosine, hypoxanthine, uric acid, and allantoin in cecal contents of the three mouse cohorts analyzed by targeted metabolomics and **E)** plasma uric acid levels analyzed by enzymatic assay were expressed as box-and-whisker plots with individual data points, where the boxes indicate the median values and the interquartile ranges and the whiskers represent the minimum and maximum values. Significance was calculated by one-way ANOVA test with the Tukey post-tests and is designated as follows: ***, *p*-value of <0.001; ****, *p*-value of <0.0001.

### Transcriptional analysis identifies bacterial genes required for anaerobic growth on UA and adenine

Having established the role of PDB on lowering levels of UA systemically we sought to identify genes encoding functions necessary for anaerobic UA catabolism. Cultures of *E. bolteae* were cultivated in medium supplemented with UA or xylose plus NH_4_Cl (henceforth, “xylose”). We selected xylose for comparison, as the growth rate of *E. bolteae* on this substrate was similar to that of UA, with doubling times of 2.6 h for xylose and 4.6 h for UA, (**Suppl. Fig. 7**). For both substrates log-phase cells were harvested and libraries subjected to sequencing (**Suppl. Fig. 7**). We obtained ~3.6 ×10^7^ reads/sample, of which 99.2% mapped to the *E. bolteae* genome. **Fig. 6** shows reads per million (RPM) normalized to gene size plotted against the relative expression level for growth on the two substrates, limited to the 3217 (of 5993) differentially expressed genes (FDR <0.01, **Suppl. Table 7**). As expected, genes encoding 30S and 50S RNA Polymerase (RNAP) subunits are indicated near the center of the figure with a slight bias (1.6-fold) towards the xylose substrate side (left), in accord with the faster growth rate on this substrate and growth rate-limiting nature of RNAP subunit expression^41^. Growth on UA promoted higher expression of 51 genes relative to xylose (cut-off >25-fold; **Fig. 6**). Eighteen of these genes encode a variety of predicted ABC/EBC/BCCT-type micronutrient transport functions including proteins likely required for metals uptake (**Suppl. Fig. 8**, filled purple circles). Also highly expressed were operons encoding a glycine cleavage system (**Suppl. Fig. 8**, filled orange circles), and a probable electron bifurcating hydrogenase (type A3; **Suppl. Fig. 8**, filled green circles), although we have not observed copious gas accumulation in these cultures and the predicted fermentation balance with UA does not require overt H2 metabolism. However, these systems have been recognized to interact with an array of substrates other than H^+^ ^42^.

**Figure 6.**
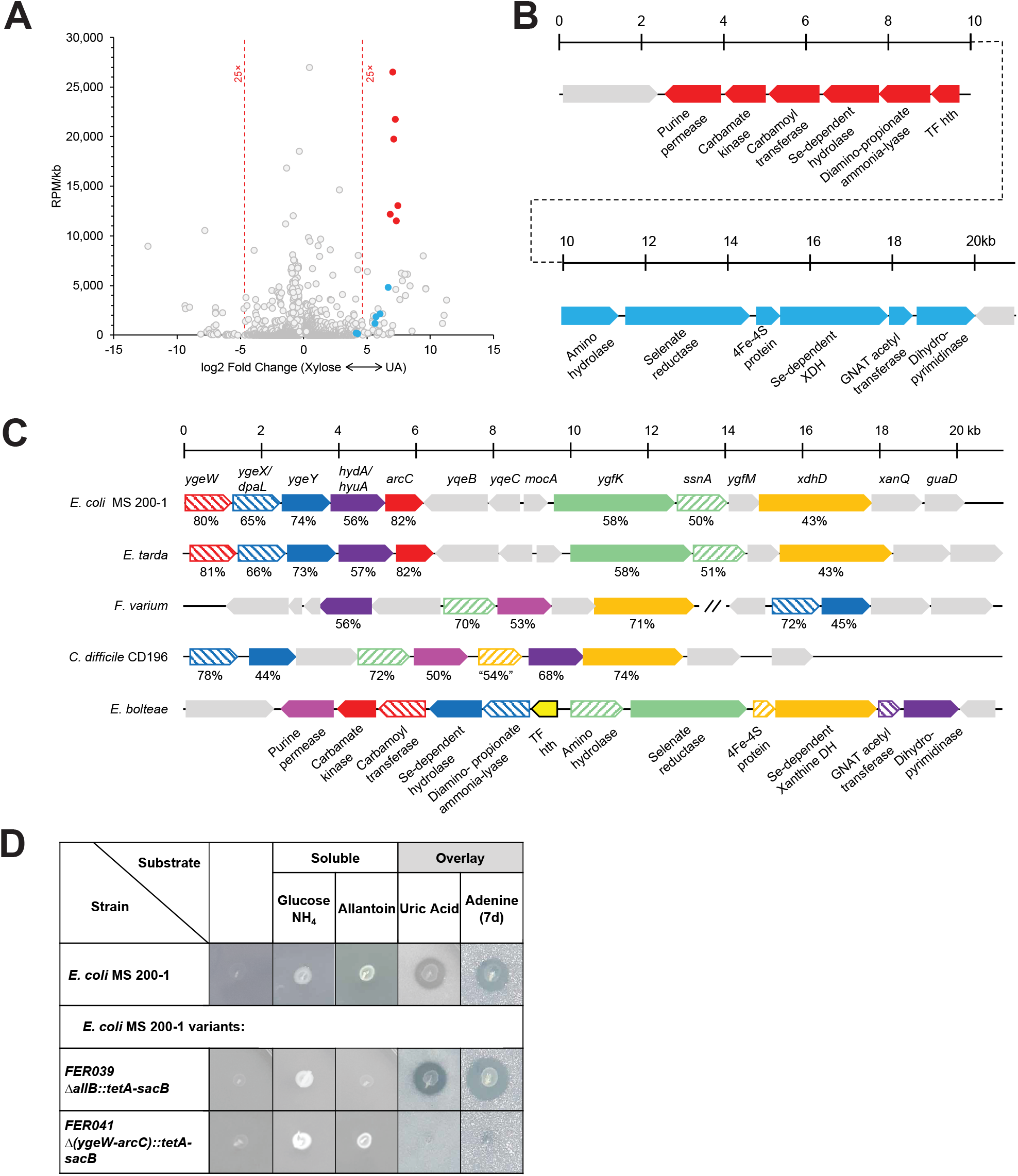
Identification of a gene cluster necessary for anaerobic bacterial growth on uric acid. **A)** Comparison of transcriptional profiles for *Enterocloster bolteae* showing differentially-expressed genes (FDR <0.01) and reads per million (RPM)/gene size (kb) for cultures grown on xylose + NH_4_Cl (“Xylose,” upregulated genes to the left) or uric acid (upregulated genes to the right), highlighting genes encoding two adjacent predicted operons encoding functions likely necessary for uric acid metabolism (blue and red circles). Descriptions of additional upregulated genes including those encoding micronutrient transport, one glycine cleavage system and a probable bifurcating hydrogenase system as well as genes upregulated during growth on xylose are described in Suppl. Fig. 8. **B)** Diagram of the adjacent upregulated predicted operons, including genes CGC65_RS20560-RS20625, color-matched with the filled circles in panel (A). Further details of gene expression is presented in Supplementary Figure 8. **C)** Representative alignments of chromosomal regions from multiple organisms that anaerobically catabolize uric acid. The genetic regions from *E. bolteae* shown in panel B are compared to those from *Clostridioides difficile* CD196 (CD196_RS16070 – RS16115), *Fusobacterium varium* (C4N18_RS01955 – RS01995) and (C4N18_RS03270 – RS03290), *Edwardsiella tarda* (ETATCC_RS03325 – RS03390) and *E. coli* MS 200-1 (HMPREF9553_RS03165 - RS03225). Matched genes are color-coded, and the percent similarities of the encoded proteins are indicated. Although selected genes appear to be conserved, their organization differs in different organisms, and in the case of *F. varium* do not occur in a contiguous chromosomal region. **D)** Growth of *E. coli* MS 200-1 wild-type and deletion variants (FER039: *ΔallB::tetA-sacB,* FER041: *Δ*(*ygeW-arcC*)::*tetA-sacB*) on plates containing glucose, allantoin, uric acid, or adenine.

Two adjacent and divergently-oriented putative operons, each encoding 6 genes, which combined amounted to 12% of all RNA-Seq reads in *E. bolteae* grown on UA. These highly upregulated genes are indicated by the filled blue and red circles with their corresponding operons and putative gene products diagrammed (**Fig. 6A, B**). Notably, the encoded proteins are predicted to catalyze numerous C-N cleavage and Se-dependent hydrolytic reactions. Also indicated are a purine permease, likely specific for UA uptake based on the conservation of residues S99 and S314 (T100 and S317 in *E. coli* UacT)^43^ and a knotted carbamoyl transferase/carbamate kinase, presumably required for ATP synthesis.

Alignments of conserved chromosomal regions of purine fermenting organisms illustrated conservation of five genes (*E. coli* nomenclature: *dpaL, hydA*, *ssnA*, *ygeY*, *xdhD*) among all taxa, although not with a conserved organization nor exclusively present in a contiguous genomic region across phyla (**Fig. 6C**). A variant of *E. coli* MS 200-1 bearing a deletion of the *ygeW*-*dpaL*-*ygeY*-*hydA*-*arcC* operon grew as the wild-type strain in medium supplied with glucose or allantoin but was unable to grow anaerobically using UA or adenine as the carbon and energy source. Conversely, a variant of *E. coli* MS 200-1 bearing a deletion of *allB*, encoding the enzyme catalyzing the first step of anaerobic allantoin metabolism^44^, was unable to utilize allantoin but retained the ability to catabolize UA and adenine, indicating distinct mechanisms of allantoin and UA / adenine metabolism in this organism (**Fig. 6D**).

### Detection of purine degradation genes in bacterial genomes and transplanted mice

Having identified genes required for anaerobic purine metabolism we then sought to identify bacterial taxa containing these genes. We performed BLASTP of NCBI RefSeq Genome Database (refseq_genomes) using parameters described in Methods. We detected 230 non-redundant bacterial taxa that had the five genes reliably detected among all experimentally confirmed purine-degrading taxa (*dpaL*, *hydA*, *ssnA*, *ygeY*, *xdhD*). These potential UA degraders included bacterial taxa belonging to Actinobacteria, Firmicutes, Proteobacteria, Fusobacteria and Spirochaetes (**Suppl. Table 8, Suppl. Fig. 9**).

Lastly we assessed the abundance of these genes in the cecum of *ApoE* gnotobiotic mice (**Fig. 1, Fig. 7A**) and correlated their abundance with levels of purine-related metabolites quantified in their cecum (**Suppl. Table 9**). Remarkably, we found that cecal levels of several purines including deoxyxanthosine, xanthosine and UA were negatively associated with the abundance of genes involved in anaerobic-purine degradation (**Fig. 7B**). Altogether these results highlight the potential of these genes as biomarkers for purine breakdown in the gut. Understanding how to manipulate the representation and function of purine-consuming species in the intestinal microbiota could potentially lead to novel means for preventing or treating hyperuricemia.

**Figure 7.**
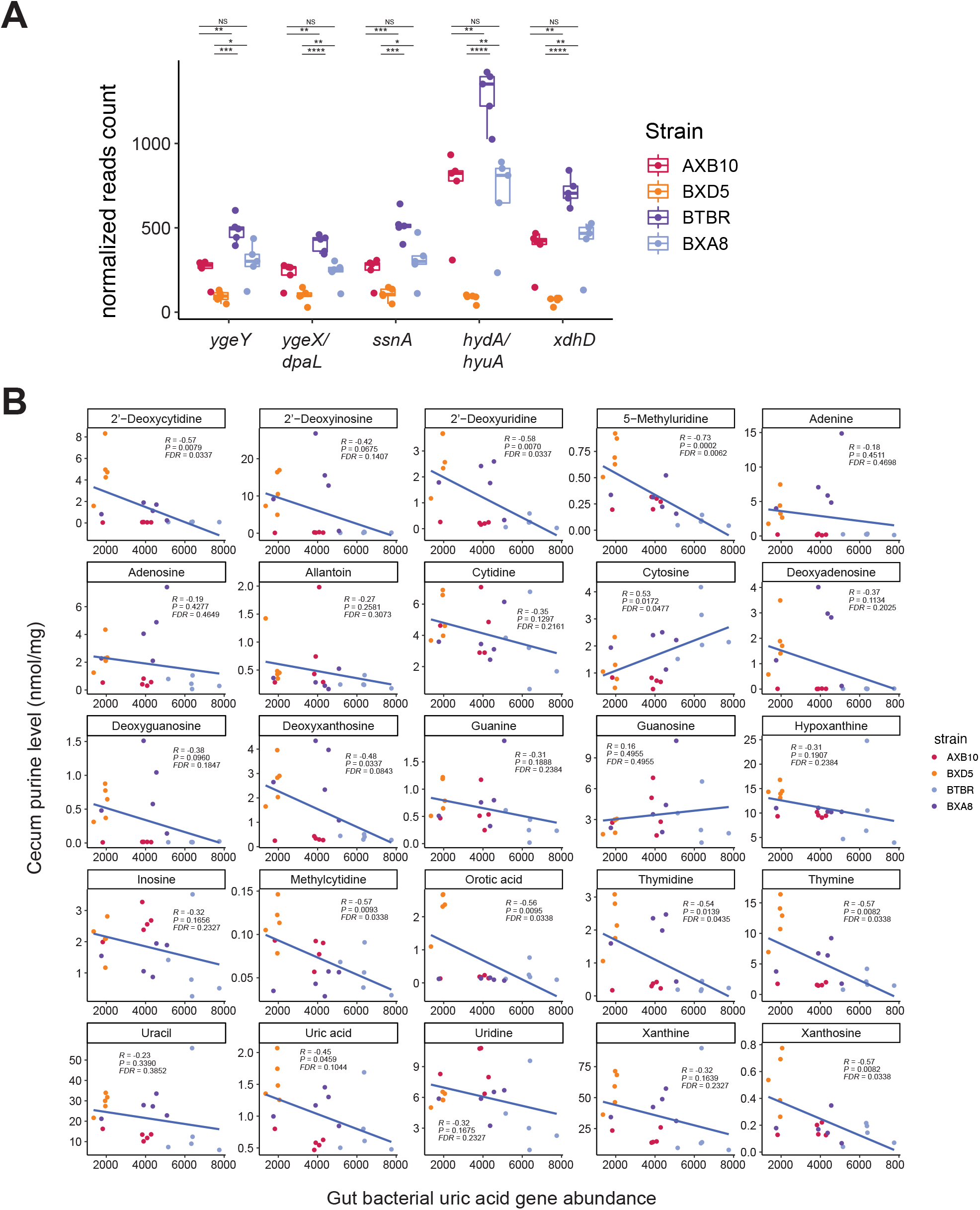
Correlation between cecal purines and gut bacterial genes involved in uric acid degradation in transplanted *ApoE* KO mice. **A)** Detection of purine degradation genes in microbiomes from transplanted mice. Abundance of genes involved in anaerobic purine degradation in gut metagenomes from gnotobiotic mice transplanted with cecal contents from strains with disparate atherosclerosis phenotypes (see Fig. 1). Differences between groups were evaluated using unpaired two-tailed Welch’s t-test. **B)** Correlation between abundance of purine-related metabolites and bacterial functions was performed using Spearman correlation.

## Discussion

In this study we sought to identify microbially-regulated metabolites involved in atherosclerosis progression. Our initial gnotobiotic mouse transplant studies revealed that levels of several purines, including UA were influenced by gut microbes and associated with atherosclerosis burden. Since hyperuricemia has been implicated in several human diseases, including cardiovascular disease, and its prevalence has been increasing in developed countries, these initial results motivated a deeper exploration into the role of the gut microbiome on purine homeostasis. These efforts led us to identify bacteria able to breakdown purines anaerobically, and to uncover a cluster of bacterial genes required for anaerobic catabolism of these substrates. We also demonstrated that taxa encoding these functions lowered circulating UA levels in mice. Altogether this work has implications beyond atherosclerosis: it provides new insights into how gut bacterial metabolism may influence UA in the circulation and suggests that microbes able to catabolize purines are important drivers of host purine homeostasis both locally (i.e., gut) and systemically.

UA is the end product of purine metabolism in humans. Nearly two-thirds of purines in the body are endogenously produced while the remainder comes from diet. Dietary purines and nucleotides are transformed to nucleosides and bases by endonucleases, phosphodiesterases, and nucleoside phosphorylases and assimilated by the host. The absorption of nucleosides through both concentrative nucleoside transporters takes place mainly in the duodenum. There is also evidence that absorption could occur in the large intestine. Absorbed purines can be used by enterocytes or colonocytes or be degraded to UA. In most mammals, UA can be further metabolized to allantoin, but in hominoids the presence of an uricase gene bearing multiple mutations and premature stop codons results in higher levels of UA^45^. Since UA is relatively insoluble, humans are susceptible to diseases resulting from precipitation of UA including gout and kidney stones. In fact, the prevalence of hyperuricemia in the U.S. is ~20% among adults and has been steadily increasing in recent decades^46,47^. This increase may be related to the prevalence of a high-purine diets, fructose beverages (known to increase UA levels), and alcohol consumption^48^. High UA levels have been previously associated with hypertension^49^, chronic kidney diseases^50^, metabolic syndrome^51^, atherosclerosis^52^ and adverse cardiovascular outcomes^53^. Our results in gnotobiotic *ApoE* KO mice (**Fig. 1**) and humans (**Fig. 2**) support this notion and suggest that levels of UA within the soluble range may contribute to disease progression. This is consistent with a recent mechanistic study suggesting that soluble UA activates the NLRP3 (NLR Family Pyrin Domain Containing 3) inflammasome^22^. Furthermore, our work extends these observations to suggest that gut bacterial catabolism of purines and UA may represent a novel mechanism by which the gut microbiome influences atherosclerosis. However, there is conflicting evidence regarding the benefits of UA-reducing strategies for treating patients with cardiovascular disease^24,25,54,55^ and more work is needed to clarify the contribution of hyperuricemia to atherosclerosis in humans.

The kidneys play a critical role in maintaining plasma UA levels through complex transport systems that mediate both reabsorption and secretion of UA^56^. Intestinal secretion is a substantial contributor to extra-renal elimination of UA, accounting for about one-third of total elimination of UA^36,37^. Our results showing that purine degrading bacteria lower the abundance of several purines in the intestine (**Fig. 5**) suggest that these organisms may lower circulating UA levels by decreasing the burden of purines bioavailable to the host. Alternatively, the increased levels of UA detected in the cecum of mice colonized with PDB (**Fig. 5**) may suggest that in the presence of these organisms, consumption of certain purines may trigger more secretion of UA into the intestine. More studies are needed to clarify how PDB influences global levels of purines and UA. Given the differential capabilities and modulation of purine consumption among the examined PDB, this may vary widely depending on the specific PDB colonizing an individual.

Anaerobic purine utilization by bacteria was first described over 100 years ago, yet relatively few species have been identified, these isolates were obtained from environmental sources and are obligate purinolytic Firmicutes. The biochemistry of their purine catabolic pathway was delineated prior to the advent of genetic manipulations and only in the past decade have their genomes been sequenced although the genetic elements encoding the process remain undefined^57,58^. More nutritionally-diverse anaerobic purine-utilizing organisms including Firmicutes and Proteobacteria isolated from termite intestines metabolize purines and evidently recycle purine nitrogen in the nitrogen-limited host diet, but further biochemical and genetic analyses of these organisms have not been published^59^. Operons encoding proposed mechanisms of anaerobic purine degradation have been identified through *in silico* analyses^60,61^, through overexpression of individual genes^62,63^ or by using cultures and cell suspensions cultivated in complex media containing additional carbon and energy sources^38^. While these efforts identified the appropriate genomic regions encoding functions necessary for anaerobic purine catabolism, the appropriate conditions allowing reproducible growth were not formulated and a full understanding of the encoded metabolism has remained elusive.

We asked to what extent this UA could serve as a source of carbon and energy for gut bacteria, and to what extent the gut microbiota composition might affect host systemic purine concentrations. We tested 34 human gut isolates from six phyla cognizant of a) the demonstrated requirements of trace elements (Mo, Se) for metabolic function^64^ and b) the well known phenomenon of catabolite repression initially reported for the *yge* operon of *E. coli* and likely present in other bacteria^65^. We identified representatives for three different phyla, primarily belonging to the Firmicutes and Proteobacteria, that readily degraded UA. A limited subset of organisms was shown to utilize adenine and, independently of the ability to use UA or adenine, some organisms including strains of *E. coli* grew anaerobically in medium supplied with allantoin (**Fig. 4, Suppl. Fig. 4**), in contrast to reports that this compound only serves as a source of nitrogen^44^. However, these properties were not consistently present in any taxonomic group and even strain differences between species were identified.

In multiple organisms the ability to utilize UA was influenced by the presence of alternative carbon sources and the abundance of metals, including Se and Mo (**Suppl. Fig. 5**). However, the effects of metals were not evident in *C. difficile* and *E. bolteae*, likely reflecting the high expression of multiple metal uptake systems identified in the RNA-Seq data for *E. bolteae* (**Suppl. Fig. 8**) and the requirement for multiple metal-limited transfers to affect mineral limitation in other purinolytic Firmicutes^64,66,67^. Whether this differential effect of trace minerals perturbs purine utilization in the gut remains to be seen. The notion that dietary Se might affect the metabolism of purines by human gut microbiota has been suggested^61,68^ but the *in vitro* results shown here indicate that any requirement could depend upon microbiota composition: necessary if purine utilizers mainly are members of the Proteobacteria but perhaps of less importance if Firmicutes predominate. Altogether, these results highlight the multiple layers that regulate purine utilization in bacteria, suggest that phylogeny is a poor predictor of microbial purine utilization and underscore the need for assessments beyond genomics when making predictions about metabolism.

Lastly, it is important to note that while the notion of using bacteria able to degrade purines might be an appealing strategy to lower pro-inflammatory UA, more work is needed to fully understand the consequences to the host. For example, the intestinal epithelium is the most vigorously self-renewing tissue of adult mammals, which imposes a high demand of nucleotides that are needed for proliferation and energy^69^. Adenine–a purine consumed by several taxa in our study (**Fig. 4**)–is a precursor of nucleic acids in intestinal cells unable to synthesize purines *de novo*^70^. Furthermore, recent work from the Colgan group showed that gut bacteria are a major source of purines that are used for nucleotide generation by intestinal mucosa. Importantly, supplementation of purines directly through bacterial colonization improved intestinal epithelial cell wound healing and barrier restitution capabilities^71,72^. This work also suggested that purines play essential roles for colonic epithelial proliferation, energy balance, and mucin barrier integrity^71,72^. Adenine also inhibits TNF (Tissue Necrosis Factor)-α signaling in intestinal epithelial cells and reduces mucosal inflammation in a dextran sodium sulfate-induced colitis mouse model^73^. Consistent with these results, a recent study identified intestinal purine starvation associated with irritable bowel syndrome^74^. Our results showing that purine degrading bacteria can breakdown adenine and lower hypoxanthine levels in the intestine, and that the abundance of genes encoding for key proteins in anaerobic purine degradation is associated with lower purines in the gut suggests that the influence of these organisms on purine availability to intestinal epithelial cells and barrier function needs to be carefully examined, especially considering that many of the taxa identified as anaerobic purine degraders, including *E. coli*, *E. bolteae*, *F. varium*, and *C. difficile* have been associated with disease^75^. Thus, further studies are warranted to examine the contribution of purine degradation to the fitness of these taxa and their health effects.

In summary, the work presented here shows that anaerobic purine utilization is widespread among gut dwelling bacteria and suggests that microbial purine degraders are important modulators of host purine homeostasis in the gut and of UA levels in circulation. Studies are needed to dissect the contribution of aerobic *vs.* anaerobic purine consuming pathways to the purine economy, gut ecology and health conditions including atherosclerosis.

## Methods

### Bacterial culture conditions

Cultures and plates were prepared in an anaerobic (ca. 75% N_2_/20% CO_2_/5% H_2_) chamber (Coy Laboratory Products, Grass Lake, MI). Media formulations and preparations are detailed in Supplementary material. Cultures were routinely grown in anaerobic septum-stoppered “Hungate tubes” (Chemglass Life Sciences, Vineland, NJ) containing a rich, well-buffered (pH 7) media, typically “Mega Medium“^76^ supplemented with maltose (0.9 g/l), cellobiose (0.86 g/l), fructose (0.46 g/l) and NaHCO_3_ (1.68 g/l) although most clostridial strains were reliably recovered from 20% glycerol freezer stocks using medium 11E. Freshly-prepared cultures were combined at roughly equivalent levels (normalized to OD600) for the purpose of colonizing germ free mice. Growth on purines utilized medium 23B, which contains 0.1% yeast extract (0.05% for the RNA-Seq cultures) as the sole undefined component, with additions of carbohydrate and NH_4_Cl (25 and 10 mM, respectively) or purines (uric acid, Sigma-Aldrich cat. # U0881, 1 mg/ml; adenine, Alfa Aesar cat. # A14906, 1 mg/ml; allantoin, Alfa Aesar cat. #A15571.30, at 45 mM) as carbon and nitrogen sources. These levels of UA and adenine did not dissolve fully, and growth in tubes was monitored by observing purine disappearance as well as culture OD600, measured using a Spectronic 20D+ (ca. 1.4 cm sample path length) after the saturating purine had settled (ca. 15 minutes). Strains and primers used in this study are listed in **Suppl. Table 10**.

Assay of anaerobic purine utilization on plates employed saturating levels of UA or adenine, using bilayer plates similar to those described previously^77^. Specifically, a 25 ml base layer consisting of a 1:1 mixture of medium 26B plus 2.4% molten Bacto agar (Becton Dickinson and Co., Franklin Lakes, NJ, catalog #214010) was poured into each 100 × 15 mm petri dish and allowed to solidify. The following day, the base layer was overlaid with a 7 ml top layer medium + agar plus (per 7 ml) 84 mg UA or 96 mg adenine. As for the liquid cultures, these levels of UA and adenine were saturating and formed and opaque overlay. Where indicated, filter-sterilized additions of concentrated stock solutions (e.g., glucose, fructose, NH_4_Cl, formate) were added to indicated levels in both base and overlay medium layers. Plates were allowed to dry for two days, then spotted with 4 μl of an overnight culture grown in rich medium (CMM or 11E) and incubated anaerobically at 37°C for 2 (UA, soluble substrates) or 7 (adenine) days, unless otherwise specified. Cultures that utilized purines both grew on the medium and formed zones of clearing as the saturating purine was depleted. Attempts to prepare overlay plates containing xanthine or guanine at levels useful to support growth did not show clear zones of purine utilization. Similarly, monolayer plates containing UA or adenine resulted in relatively indistinct zones of clearing relative to bilayer plates, often with formation of a dense ammonium urate precipitate especially after storate at 4°C. Five cultures were spotted/plate--a *E. bolteae* positive control plus 4 test strains.

### Test of trace mineral requirements for growth on UA

Tests utilized the standard phosphate-buffered basal medium formulation (“26B” with added 100 nM Mn, Ni, Zn, 50 nM Co, W) plus 2 out of 3 of the following: 2.5 μM Fe, 5 μM Mo, 0.5 μM Se (specific mineral compounds are listed in **Supplementary Methods**). No attempt was made to rigorously exclude the individual metal being tested, and it is likely that other medium components—particularly phosphate buffer, cysteine and yeast extract—supplied trace levels; therefore, evidence of mineral requirements indicates a substantial demand.

### HPLC analysis

Analyses were performed using a Shimadzu (Columbia, MD) system comprised of a CBM-40 controller, LC-40D pumps, SIL-40C autosampler, CTO-40C column oven, and SPD-M40 diode array detector. Samples were applied to a Phenomenx (Torrace, CA) Luna Omega 5μm Polar C18 LC column (250 × 4.6 mm) maintained at 25°C with a 0.5 ml/min gradient composed of (A) 100 mM K_x_H_x_PO_4_ pH 2.4 and (B) A with 40% acetonitrile as follows (Time in minutes, %A, %B): 0, 99, 1 / 15, 96, 4 / 25, 25, 75 / 35, 25, 75 / 36, 99, 1 / 50, 99,1. Under these conditions formate, lactate and acetate eluted at 7.1, 10.1, and 10.6 minutes and 1 mM concentrations were readily detected at 205 nm. UA eluted at 17.4 minutes and 10 μM was detected at 284 nm. As some samples contained saturating substrate levels, for UA analyses all samples were vigorously mixed and immediately diluted 40-fold in phosphate buffered saline to allow full UA dissolution prior to analysis.

### Headspace Gas Chromatography

Analyses of short-chain fatty acids, and ethanol were performed using a Shimadzu headspace GC/FID as previously described^78^.

### RNA isolation and sequencing

10-ml cultures were grown in triplicate in tubes containing medium 23B (with 0.05% yeast extract) plus a) 25 mM xylose + 10 mM NH_4_Cl or b) 12 mg UA and harvested at OD600 = 0.25 by plunging the tubes into an ice water slurry, then pelleting cells prior to storage at −80°C. RNA was isolated using the Monarch Total RNA Miniprep kit from New England Biolabs (Ipswich, MA), with yields of 2000 – 3000 ng RNA per sample, assayed by dye binding (Qubit, Invitrogen / Thermo Fisher Scientific, Waltham, MA). Samples in were submitted to the Microbial Genome Sequencing Center (MiGS, Pittsburgh, PA) where Illumina Stranded RNA library preparation with RiboZero Plus rRNA depletion was performed, followed by Illumina sequencing [paired-end reads (2×51bp)].

### Mouse experimental design

We conducted three animal experiments as follows. i) At UCLA six-week-old female AXB10/PgnJ, BTBR T+tf/J, BXD5/TyJ, and BXA8/PgnJ mice were fed a Western diet (Research Diets D10042101) for 4 weeks, and cecal contents were collected. Cecal samples were shipped to the University of Wisconsin-Madison for microbiota transplant. Ten-week-old GF female *ApoE* KO mice (C57BL/6J background) fed a standard chow (LabDiet 5021; LabDiet, St Louis, MO) were inoculated by oral gavage with 0.2 mL of resuspended cecal slurry from the HMDP strains. Mice were switched to a standard chow diet supplemented with 0.2% cholesterol (TD.07798; Envigo, Madison, WI). Mice were then euthanized at 18 weeks of age after 4h fasting and collected tissues. ii) Cecal and plasma samples were collected from eighteen-week-old GF or conventionally-raised *ApoE* KO mice fed the 0.2% cholesterol-supplemented diet (TD.07798) for 8 weeks. iii) Three groups of adult gnotobiotic C57BL/6J mice were tested for purine metabolism: a) the “core” community which included seven species that do not degrade purines *in vitro*, *Bacteroides caccae*, *Bifidobacterium dentium*, *Blautia hansenii*, *Bacteroides thetaiotaomicron*, *Coprococcus comes*, *Mitsuokella multacida*, and *Ruminococcus torques*; b) the “core plus purine-degrading bacteria (PDB)” community that added three PDB to the “core” community mixture including *E. bolteae*, *H. hathewayi,* and *E. coli*; and c) the last group remained germ-free. The three groups of mice were fed a standard chow (LabDiet 5021). Cecal and plasma samples were collected 4 weeks after the colonization. The animal works were conducted according to relevant national and international guidelines and was approved by the UCLA Animal Research Committee, the UCLA IACUC, or the University of Wisconsin-Madison Animal Care and Use Committee.

### Gnotobiotic husbandry

All GF C57BL/6J and *ApoE* KO mice were maintained in a controlled environment in either plastic flexible film gnotobiotic isolators or individually ventilated cages under a strict 12 h light cycle and received sterilized water and standard chow (LabDiet 5021) *ad libitum* unless otherwise stated. Sterility of GF animals was assessed by routine PCR testing (16S rRNA gene) and by incubating freshly collected fecal samples under aerobic and anaerobic conditions using standard microbiology methods.

### Atherosclerotic lesion assessments

Atherosclerotic lesions were assessed as previously described^79^. Briefly, mice were anesthetized and the aorta was perfused with PBS. To assess the atherosclerotic lesion size at the aortic sinus, the samples were cut in the ascending aorta, and the proximal samples containing the aortic sinus were embedded in OCT compounds (Tissue-Tek; Sakura Finetek, Tokyo, Japan). Five consecutive sections (10 μm thickness) taken at 100 μm intervals (i.e. 50, 150, 250, 350, and 450 μm from a bottom of the aortic sinus) were collected from each mouse and stained with Oil Red O. Plaque area and Oil Red O-positive area were measured using Image J software (National Institutes of Health, Bethesda, MD). The volume of atherosclerosis in the aortic sinus was expressed as mean size of the 5 sections for each mouse.

Immunohistochemistry was performed on formalin-fixed cryosections of mouse aortic roots using antibodies to identify macrophages (MOMA-2, 1:50; ab33451, Abcam, Cambridge, MA), followed by detection with biotinylated secondary antibodies (1:400; ab6733, Abcam) and streptavidin-horseradish peroxidase (1:500; P0397, Dako, Carpinteria, CA). Smooth muscle cells were identified by immunostaining with fluorescein isothiocyanate (FITC)-conjugated primary antibody against α-smooth muscle actin (1:100; clone 1A4, Sigma), followed by anti-FITC biotin-conjugated secondary antibody (1:400; clone FL-D6, Sigma). Negative controls were prepared with substitution with an isotype control antibody. Staining with Masson’s trichrome was used to delineate the fibrous area according to the manufacturer’s instructions (ab150686, Abcam). Stained sections were digitally captured, and the percentage of the stained area (the stained area per total atherosclerotic lesion area) was calculated.

### Plasma biochemical analysis

Blood samples were collected by cardiac puncture into EDTA-rinsed syringes under anesthesia using isoflurane. Plasma was acquired by centrifugation and stored at −80°C until measurement. The levels of triglycerides, total cholesterol, and high-density lipoprotein cholesterol were measured with commercially available kits from Wako Chemicals (Richmond, VA). Plasma LPS levels were quantitated with the QCL-1000 Endpoint Chromogenic LAL Assay (Lonza, Basel, Switzerland). Plasma UA levels were determined by the Vistro DT60 II Analyzer at the University of Massachusetts Medical School MMPC (National Mouse Metabolic Phenotyping Center).

### uHPLC-MS/MS analysis of metabolites

Plasma samples were prepared for analysis by precipitating proteins with 4 volumes of ice-cold methanol spiked with 2.5 μM deuterium-labeled choline and TMAO internal standards. Samples were centrifuged at 18,213 x *g* at 4°C for 3 min. The recovered supernatants were diluted 1:1 in uHPLC-grade water prior to screening. Identification and quantification of choline and TMAO was performed using a uHPLC (Dionex 3000) coupled to a high-resolution mass spectrometer (Thermo Scientific Q Exactive). Liquid chromatography separation was achieved on a Dikma Bio-Bond C_4_ column (150 mm by 2.1 mm; 3-μm particle size) using a 7-min isocratic gradient (50:50 methanol-water, 5 mM ammonium formate, and 0.1% formic acid). A heated electrospray ionization interface, working in positive mode, was used to direct column eluent to the mass spectrometer. Quantitation of TMAO and D_9_-TMAO was performed via targeted MS/MS using the following paired masses of parent ions and fragments: TMAO (76.0762 and 58.0659) and D_9_-TMAO (85.1318 and 68.1301). Quantitation of choline and d_9_-choline was performed in full-MS scan mode by monitoring their exact masses: 104.1075 and 113.1631, respectively.

### GC-MS of short chain fatty acid measurement

Sample preparation was based on a previously described procedure^10^. Cecal contents were weighed in 4mL polytetrafluoroethylene (PTFE) screw cap vials and 10 μL of a mixture of internal standards (20 mM of acetic acid-D4, Sigma-Aldrich #233315; propionic acid-D6, Sigma-Aldrich #490644; and butyric acid-D7, CDN isotopes #D-171) was subsequently added to each vial, followed by 20 μL of 33% HCl and 1 mL diethyl ether. For plasma samples, 50 μL of each sample, 1.25 μL of the internal standard mix, 5 μL of 33% HCl, and 0.75 mL of diethyl ether were mixed. The mixture was vortexed vigorously for 3 min and then centrifuged (4,000 x *g*, 10 min). The upper organic layer was transferred to another vial and a second diethyl ether extraction was performed. After combining the two ether extracts, a 60 μL aliquot was removed, combined with 2 μL *N-tert*-butyldimethylsilyl-*N*-methyltrifluoroacetamide (MTBSTFA, Sigma-Aldrich #394882) in a GC auto-sampler vial with a 200 μL glass insert, and incubated for 2 h at room temperature. Derivatized samples (1 mL) were injected onto an Agilent 7890B/5977A GC/MSD instrument with a Agilent DB1-ms 0.25 mm x 60 m column with 0.25 μm bonded phase. A discontinuous oven program was used starting at 40°C for 2.25 min, then ramping at 20°C/min to 200°C, then ramping at 1 00°C/min to 300°C and holding for 7 min. The total run time was 18.25 minutes. Linear column flow was maintained at 1.26mL/min. The inlet temperature was set to 250°C with an injection split ratio of 15:1. Quantitation was performed using selected ion monitoring (SIM) acquisition mode and metabolites were compared to relevant labeled internal standards using Agilent Mass Hunter v. Acquisition B.07.02.1938. The m/z of monitored ions are as follows: 117 (acetic acid), 120 (acetic acid-D4), 131 (propionic acid), 136 (propionic acid-D6), 145 (butyric acid), and 151 (butyric acid-D7). Concentrations were normalized to mg of cecal contents.

### Ultrahigh Performance Liquid Chromatography-Tandem Mass Spectroscopy (UPLC-MS/MS) for untargeted metabolome for plasma samples

Untargeted mass spectrometry data were collected at Metabolon Inc (Durham, NC). Plasma samples were prepared using the automated MicroLab STAR system (Hamilton Company, Bonaduz, Switzerland). To remove protein, dissociate small molecules bound to protein or trapped in the precipitated protein matrix, and to recover chemically diverse metabolites, proteins were precipitated with methanol under vigorous shaking for 2 min (Glen Mills GenoGrinder 2000) followed by centrifugation. The resulting extract was divided into five fractions: two for analysis by two separate reverse phase (RP)/UPLC-MS/MS methods with positive ion mode electrospray ionization (ESI), one for analysis by RP/UPLC-MS/MS with negative ion mode ESI, one for analysis by HILIC/UPLC-MS/MS with negative ion mode ESI, and one sample was reserved for backup. Samples were placed briefly on a TurboVap (Zymark) to remove the organic solvent. The sample extracts were stored overnight under nitrogen before preparation for analysis.

All methods utilized a Waters ACQUITY ultra-performance liquid chromatography (UPLC) and a Thermo Scientific Q-Exactive high resolution/accurate mass spectrometer interfaced with a heated electrospray ionization (HESI-II) source and Orbitrap mass analyzer operated at 35,000 mass resolution. The sample extract was dried then reconstituted in solvents compatible to each of the four methods. Each reconstitution solvent contained a series of standards at fixed concentrations to ensure injection and chromatographic consistency. One aliquot was analyzed using acidic positive ion conditions, chromatographically optimized for more hydrophilic compounds. In this method, the extract was gradient eluted from a C18 column (Waters UPLC BEH C18–2.1×100 mm, 1.7 μm) using water and methanol, containing 0.05% perfluoropentanoic acid (PFPA) and 0.1% formic acid (FA). Another aliquot was also analyzed using acidic positive ion conditions, however it was chromatographically optimized for more hydrophobic compounds. In this method, the extract was gradient eluted from the same aforementioned C18 column using methanol, acetonitrile, water, 0.05% PFPA and 0.01% FA and was operated at an overall higher organic content. Another aliquot was analyzed using basic negative ion optimized conditions using a separate dedicated C18 column. The basic extracts were gradient eluted from the column using methanol and water, amended with 6.5mM ammonium bicarbonate at pH 8. The fourth aliquot was analyzed via negative ionization following elution from a HILIC column (Waters UPLC BEH Amide 2.1 × 150 mm, 1.7 μm) using a gradient consisting of water and acetonitrile with 10mM ammonium formate, pH 10.8. Compounds were identified by comparison to library entries based upon retention time/index, mass to charge ratio (m/z) and chromatographic data, and peaks were quantified using area-under-the curve.

### Targeted purine metabolite quantitation by LC-MRM/MS

Purines mass spectrometry data were collected at TMIC (The Metabolomics Innovation Centre, UVic Genome BC Proteomics Centre, Victoria, Canada). An internal standard (IS) solution containing 13C- and/or 15N-labeling AMP, ATP, GMP, GTP, UMP, UTP, xanthine, guanine and adenine, hypothanxine, guanosine and adenosine was prepared in 80% methanol (for cecal samples) or 10% methanol (for plasma samples). Serially diluted standard solutions containing standard substances of the targeted nucleotides, nucleosides and nucleobases were prepared in a concentration range of 0.00001 to 50 nmol/mL in 80% methanol (for cecal samples) or 0.00001 to 10 nmol/mL in 10% methanol (for plasma samples). The cecal samples were lyophilized to dryness. The powder of each sample was weighted to an Eppendorf tube. 80% methanol at 20 μL per mg of powder was added. The samples were homogenized on a mill mixer at 30 Hz for 2 min, three times, with the aid of 2 metal beads, followed by sonication in an ice-water bath for 5 min. The tubes were centrifuged at 21,000 g and 5 °C for 20 min. 100 μL of the clear supernatant of each sample or 100 μL of each standard solution was in turn mixed with an equal volume of the IS solution, 100 μL of water and 120 μL of dichloromethane. The mixtures were vortex-mixed at 3,000 rpm for 2 min and subsequently centrifuged at 21,000 g for 5 min. 120 μL of the clear supernatant was collected and then dried at 30 °C under a nitrogen gas flow. 20uL of each plasma, once thawed on ice, was mixed with 200 μL of the IS solution and 780 uL of methanol. After vortex mixing at 3000 rpm for 1 min and sonication in an ice-water batch for 2 min, the sample was centrifuged at 21,000 g and and 5 °C for 15 min. 500 μL of the clear supernatant of each sample was transferred to another tube, 100 μL of water and 400 μL of dichloromethane was added. The mixture was vortex mixed at 3000 rpm for 1 min and then centrifuged. The upper aqueous phase of each sample was taken out to a LC micro-vial and then dried under a nitrogen gas flow. For both cecal and plasma samples, the residues were reconstituted in 100 μL of 10% methanol. 10-μL aliquots of the sample solutions and the standard solutions were injected into a C18 LC column (2.1*100 mm, 1.8 μm) to run UPLC-MRM/MS on a Waters Acquity UPLC system coupled to a Sciex QTRAP 6500 Plus mass spectrometer operated in the negative-ion mode for detection of nucleotides. The mobile phase was a tributylamine acetate buffer and acetonitrile for binary gradient elution (5% to 35% acetonitrile over 25 min), at 0.25 mL/min and 40 °C. For quantitation of nucleosides and nucleobases, 10-μL aliquots of the sample solutions and the standard solutions were injected into a polar C18 UPLC column (2.1×100 mm, 1.6 μm) to run UPLC-MRM/MS on the same LC-MS instrument operated in the positive-ion mode. The mobile phase was a 0.1% HFBA solution and acetonitrile for binary-solvent gradient elution (0% to 28% acetonitrile in 14 min), at 0.30 mL/min and 40 °C. Concentrations of detected analytes were calculated with internal-standard calibration by interpolating the constructed linear-regression curves of individual compounds, with the analyte-to-internal standard peak area ratios measured from the sample solutions.

### DNA Extraction from cecal and fecal samples

DNA was extracted from samples according to published bead-beating procedures^10^. In short, fecal or cecal samples were resuspended in a solution containing 500 μl of 2X extraction buffer [200 mM Tris (pH 8.0), 200 mM NaCl, 20 mM EDTA], 210 μl of 20% SDS, 500 μl phenol:chloroform:isoamyl alcohol (pH 7.9, 25:24:1) and 500 μl of 0.1-mm diameter zirconia/silica beads. Cells were mechanically disrupted using a bead beater (BioSpec Products, Barlesville, OK) for 3 min at room temperature. The aqueous layer was removed and DNA precipitated using 600μl isopropanol and 60μl 3M Na-acetate. Pellets were dried with ethanol and resuspended in TE. NucleoSpin Gel and PCR Clean-up Kit (Macherey-Nagel, Bethlehem, PA) was used to remove contaminants. Isolated DNA was stored at −80°C until downstream processing.

### Metagenomic shotgun DNA sequencing

DNA was extracted from cecal contents of individual mice using previously described methods^80,81^. Following DNA extraction, Illumina paired end libraries were constructed using a previously described protocol^40^, with a modification of gel selecting DNA fragments at ∼450 bp in length. Paired end (PE) reads (2 x 125) were generated using HiSeq 2500 platform.

### Metagenomic reads processing

Raw reads were preprocessed using Fastx Toolkit (ver. 0.0.13): (1) for demultiplexing raw samples, fastx_barcode_splitter.pl, with –partial 2, mismatch 2 was used; (2) when more than one forward and reverse read file existed for a single sample (due to being run on more than one lane, more than one platform, or at more than one time), read files were concatenated into one forward and one reverse read file; (3) barcodes were trimmed to form reads (fastx_trimmer -f 9 -Q 33); (4) and reads were trimmed to remove low quality sequences (fastq_quality_trimmer -t 20 -l 30 -Q33). Following trimming, unpaired reads were eliminated from the analysis using custom Python scripts. To identify and eliminate host sequences, reads were aligned against the mouse genome (mm10/GRCm38) using bowtie2^82^ (ver. 2.3.4) with default settings and microbial DNA reads that did not align to the mouse genome were identified using samtools (ver. 1.3) (samtools view -b -f 4 -f 8).

### Microbiome trait quantification

Quantification of microbial genes was done by aligning clean PE reads from each sample to a previous published mouse gut microbiome NR gene catalog using Bowtie2 (ver. 2.3.4) and default parameters. RSEM^83^ (ver. 1.3.1) was used to estimate microbial gene abundance. Relative abundance of microbial gene counts per million (CPM) were calculated using microbial gene expected counts divided by gene effective length then normalized by the total sum. To obtain abundance information for microbial functions, CPM of genes with the same KEGG Orthology (KO) annotation were summed together. In case there were multiple KO annotations for a single gene, we used all KO annotations. To obtain taxonomic abundance, CPM of genes with the same NCBI taxa annotation were summed together at phylum, order, class, family, and genus levels with a minimum of 10 genes in each taxon.

### COPRO-Seq analysis

Bacteria communities resulting from inoculation of GF animals were analyzed using Illumina sequencing according to the COPRO-Seq (*co*mmunity *pro*filing by *seq*uencing) method^84^. Feces were collected 4 weeks after the colonization. In short, DNA isolated from feces via bead beating was used to prepare libraries for shotgun Illumina sequencing. Five hundred nanograms of DNA from each sample was fragmented by sonication and subjected to enzymatic blunting and adenine tailing. Customized Illumina adapters containing maximally distant 8-bp barcodes were ligated to the poly (A)-tailed DNA. Gel-extracted DNA (size selection ~250 to 300bp) was amplified by PCR using primers and cycling conditions recommended by Illumina. Purified PCR products were submitted to the UW-Madison Biotechnology Center for a single end 50-bp Illumina MiSeq run. Results were processed using the software pipeline detailed by McNulty *et al*^84^.

### Human study

UA and coronary artery calcification (CAC) data obtained from pre-diabetes Swedish cohort participating in a study examining the link between the gut microbiota and type 2 diabetes^34^. UA was measured using a photometric technique on a Roche Cobas analyzer (Clinical Chemistry, Sahlgrenska University Hospital). Coronary artery calcifications (CAC) were assessed by computed tomography (CT) scanning using a dual-source CT scanner equipped with a Stellar Detector (Siemens, Somatom Definition Flash, Siemens Medical Solution, Forchheim, Germany). CAC images were obtained using electrocardiogram-gated non-contrast cardiac CT imaging at 120 kV. All non-contrast image sets were reconstructed (B35f HeartView medium CaScore) and CAC were identified and scored using the syngo.via calcium scoring software (Volume Wizard; Siemens) to obtain a CAC score according to Agatston. Previously reported microbiome data^34^ was mapped against Unified Human Gastrointestinal Genome (UHGG) v1.0 catalogue^85^ using Kraken2 v.2.1.2^86^ to examine association between UA levels and bacterial taxa. Xgboost^87^ and caret packages^88^ in R version 4.0.3 were used to select microbes associated to the serum urate levels based on regression analysis. These associations were further assessed for significance after adjustment for covariates (CACS, BMI, gender, triglycerides and HbA1c) in a mixed linear regression model.

### Inactivation of *allB* and *ygeW-arcC* in *E. coli* MS 200-1

The *allB* gene (Locus: NZ_GG773866; HMPREF9553_RS01540) and the *ygeW-arcC* operon (HMPREF9553_RS03165 - RS03180) were deleted and replaced with the *tetA-sacB* cassette amplified from T-SACK in accord with standard recombineering methods, followed by elimination of the pSIM5 helper plasmid^89^. The resulting constructs were verified by sequencing across the cassette-genome junctions. Primers used in these constructs are listed in **Suppl. Table 10**. Unfortunately *E. coli* MS 200-1 proved to be resistant to phage P1 transduction and the constructs could not be transferred to naïve recipients.

### Isolation of strain *E. coli* I-11 from a human fecal samples

This strain was isolated from a human fecal sample collected in accord with University of Wisconsin Health Science Institutional Review Board. For the isolation, a small amount (approximately 20 mg) of aseptically-sampled material was injected into a 10-ml medium 23B anaerobic culture supplemented with 10 mg of UA, then incubated at 37°C. Upon observation of growth and disappearance of the UA precipitate, the culture was transferred (1:100 dilution) into the same medium, maintained under the same conditions, this was repeated then streaked to UA bilayer plates. Colonies demonstrating UA metabolism were purified and presumptively identified by sequencing of the 16S rRNA gene. Strain I-11 was a facultative rod-shaped organism, which fermented glucose, sucrose and lactose but not cellobiose, as expected for *E. coli*^90^.

### Quantification of purine degradation genes

Purine degradation genes from bacterial genomes were predicted by BLASTP of NCBI RefSeq Genome Database (refseq_genomes) using five of *Enterocloster bolteae* UA utilization genes as query (*ygeY*, Se-dependent hydrolase; *ygeX*/*dpaL*, Diamino-propionate ammonia-lyase; *ssnA*, Amino-hydrolase; *hydA*/*hyuA*, Dihydro-pyrimidinase; *xdhD*, Se-dependent Xanthine DH). Purine degradation bacteria were defined by their genomes containing the five genes listed above with identity > 25% to *Enterocloster bolteae* UA utilization genes. We removed redundancy if two bacteria genomes have identical protein for all five purine degradation genes. To quantify the abundance of purine degradation genes in gut microbiomes of transplanted mice, we mapped metagenomic reads to identify purine degradation genes using RSEM (ver. 1.3.1)^83^. Relative abundance of microbial gene counts per million (CPM) were calculated using microbial gene expected counts divided by gene effective length then normalized by the total sum.

### Data analysis and statistical analysis

Data integration and statistical analysis were performed in R (ver. 3.6.3) or Prism 9. The data were expressed as box-and-whisker plots with individual data points, where the boxes indicate the median values and the interquartile ranges and the whiskers represent the minimum and maximum values. Significance was calculated by unpaired two-tailed Student’s t-test or one-way ANOVA with the Tukey post-tests. The correlation analysis was performed using Spearman’s correlation by R function “cor.test()”. For multiple testing, Benjamini-Hochberg FDR procedure was used to adjust p-values. Heatmap plots were performed using R package pheatmap (ver. 1.0.12).

### Data availability

The accession number for sequence data used in the study (Mouse Metagenomics data, Bacterial RNA-Seq data, Copro-Seq data, Human Metagenomics data) are PRJNA904303, PRJNA903666, and EGAS00001004480 (European Genome-Phenome Archive).

## Acknowledgements

We thank the University of Wisconsin Biotechnology Center DNA Sequencing Facility for providing sequencing and support services, the University of Wisconsin Mass Spectrometry Facility for the SCFA measurement, the Metabolomics Innovation Centre (TMIC) for purine metabolomics analysis, and Metabolon for untargeted metabolomics analysis. This work was partly supported by grants from NIH HL 144651 (F.E.R. and A.L.) and NIH HL148577 (F.E.R. and A.L.). This work was also supported by a grant from a Transatlantic Networks of Excellence Award from Foundation Leducq (17CVD01; to F.B. and F.E.R.) and Knut and Alice Wallenberg Foundation (2017.0026; to F.B.). F.B. is the Torsten Söderberg Professor in Medicine and a Wallenberg Scholar.

## Author contributions

KK and FER conceived the study. KK, and RLK performed microbiology, mouse studies and collected phenotypic and transcriptomic data. QZ and MP conducted statistical analyses. MM, AL contributed key reagents. MP, GB and FB conducted human studies. KK, RLK and FER wrote the manuscript. All authors read and approved the final manuscript.

## Figure legends

**Supplementary Figure 1. Body weight, lipid profile, and microbial metabolites from transplanted mice. A)** Body weight and epididymal fat weight collected at the end of the experiment. **B)** Total cholesterol, triglyceride, and HDL-cholesterol in plasma. **C)** Plasma lipopolysaccharide (LPS) levels. **D)** Plasma TMAO and choline levels. **E)** Cecal acetate, propionate, and butyrate levels. Data are shown as box-and-whisker plots with individual data points, where the boxes indicate the median values and the interquartile ranges and the whiskers represent the minimum and maximum values. Significance was calculated by one-way ANOVA with the Tukey post-tests and is reported as follows: *, *p*-value of <0.05; **, *p*-value of <0.01.

**Supplementary Figure 2. Metagenomic analysis of transplanted *ApoE* knockout mice. A)** Relative abundance of molecular functions (Kegg Orthology, KO) involved in purine metabolism in transplanted *ApoE* knockout mice. **B)** Spearman correlation between bacterial KO involved in purine and atherosclerosis lesion size.

**Supplementary Figure 3. Targeted purine metabolite quantitation in plasma samples from Conv and GF mice. A)** Heatmap of purine metabolites in plasma samples from Conv (n=8) and GF (n=8) mice analyzed by LC-MS/MS. **B)** PLS-DA plot based on the data derived from purine metabolites in plasma samples from Conv and GF mice. **C)** VIP plot indicating the most discriminating metabolite in descending order of importance. The colored boxes on the right indicate the relative concentrations of the corresponding metabolite in each group. Conv; Conventionally-raised, GF; germ-free, PLS-DA; Partial Least Squares Discriminant Analysis, VIP; variable importance of projection.

**Supplementary Figure 4. Screen of gut bacterial isolates for growth on purines.** Overnight bacterial cultures were spotted (4 μl) onto medium 26B agar plates, and plates containing soluble additions (NH_4_ = 10 mM NH_4_Cl, Glucose = 25 mM glucose, Allantoin = 25 mM Allantoin) or overlays containing saturating levels of uric acid (UA), Adenine, or UA plus formate (25 mM), as detailed in the Materials and Methods. Plates were incubated anaerobically at 37°C for 2 (all except with adenine overlay) or 7 days (adenine overlay). * indicates no test performed,

**Supplementary Figure 5. Effects of trace minerals and sugars on purine utilization.** Trace minerals: Plates containing 20 mM glucose + 10 mM NH_4_Cl, or bilayer uric acid overlay plates were prepared with different trace mineral compositions: i) containing all trace element additions (see Materials and Methods with 2.5 μM Fe, 5 μM Mo and 0.5 μM Se) or ii) lacking the addition of the indicated individual trace element. No attempt was made to rigorously remove the “missing” minerals, and plates were prepared with otherwise standard cysteine·HCl-reduced, phosphate-buffered medium 26B containing 0.1% yeast extract and Difco Bacto agar. Sugars: Bilayer uric acid plates were prepared as described or supplemented with filter-sterilized stocks of NH_4_Cl (to 10 mM), fructose and NH_4_Cl (to 40 and 10 mM, respectively), or glucose + NH_4_Cl (to 40 and 10 mM, respectively). Plates were spotted with 4 μl of cultures freshly-grown in rich medium and incubated as usual for 2 days at 37°C.

**Supplementary Figure 6. Targeted purine metabolite quantitation in plasma samples from GF and gnotobiotic mice. A)** Heatmap of purine metabolites in plasma samples from GF (n=5), ‘core’ (n=3) and ‘core plus PDB’ (n=5) mice analyzed by LC-MS/MS. **B)** PLS-DA plot based on the data derived from purine metabolites in plasma samples from GF, ‘core’ and ‘core plus PDB’ mice. **C)** Variable Importance Projection plot indicating the most discriminating metabolite in descending order of importance. The colored boxes on the right indicate the relative concentrations of the corresponding metabolite in each group. GF; germ-free, PDB; purine-degrading bacteria, PLS-DA; Partial Least Squares Discriminant Analysis, VIP; variable importance of projection.

**Supplementary Figure 7. Growth of *E. bolteae* cultures for transcriptional profiling.** Anaerobic Hungate tubes were prepared with medium 23B (with 0.05% yeast extract as the sole undefined component) and (in triplicate) a) 25 mM Xylose + 10 mM NH_4_Cl or b) 12 mg uric acid. Cultures were incubated without shaking (37°C) and were mixed approximately 15 minutes prior to each OD measurement, which allowed for settling of the uric acid precipitate. Cells were harvested at OD600 ~ 0.25. The recovered cell pellets were frozen and stored at −80°C prior to RNA extractions.

**Supplementary Figure 8. Comparison of transcriptional profiles for *E. bolteae* grown on xylose *vs.* uric acid.** Plot showing differentially-expressed genes (FDR <0.01) and reads per million (RPM)/gene size (kb) for *Enterocloster bolteae* grown on xylose + NH_4_Cl (upregulated genes to the left) or uric acid (upregulated genes to the right). Genes encoding 30S (yellow “x”) and 50S (red “x”) RNAP subunits are indicated near the center of the figure. Growth on xylose + NH_4_Cl elicited high expression of genes for sugar transport functions, an operon encoding xylose-utilization proteins, and alcohol dehydrogenases, the latter in accord with the accumulation of ethanol in these cultures. In addition to the two operons described in the manuscript, growth on uric acid also induced high expression of micronutrient transport functions, one glycine cleavage system, and a bifurcating hydrogenase system. Relevant genes are indicated in the right-hand panels, color-coded according to the expression plots shown on the left.

**Supplementary Figure 9. Detection of purine degradation genes in bacterial genomes.** Purine degradation genes from bacteria genomes were predicted by BLASTP of NCBI RefSeq Genome Database (refseq_genomes) using five of *Enterocloster bolteae* uric acid utilization genes as queries (*ygeY*, Se-dependent hydrolase; *ygeX*/*dpaL*, Diaminopropionate ammonia-lyase; *ssnA*, Amino-hydrolase; *hydA*/*hyuA*, Dihydro-pyrimidinase; *xdhD*, Se-dependent Xanthine DH). Heatmap indicating taxa, percent identity between *E. bolteae* and genomes containing at least five genes reliably detected among all experimentally confirmed purine-degrading taxa.

**Supplementary Table 1. Metabolites detected in plasma samples from gnotobiotic *ApoE* KO mice**

**Supplementary Table 2. Fecal bacterial taxa associated with uric acid levels in humans**

**Supplementary Table 3. Abundance of purine related metabolites in cecal contents of germ-free and conventional mice**

**Supplementary Table 4. Abundance of purine related metabolites in plasma of germ-free and conventional mice**

**Supplementary Table 5. Abundance of purine related metabolites in cecal contents from gnotobiotic mice colonized with purine-degrading bacteria**

**Supplementary Table 6. Abundance of purine related metabolites in plasma from gnotobiotic mice colonized with purine-degrading bacteria**

**Supplementary Table 7. Genes differentially expressed by *E. bolteae* grown on uric acid compared to xylose/NH_4_Cl.**

**Supplementary Table 8. Prediction of genes encoding proteins required for growth on uric acid**

**Supplementary Table 9. Purine-related metabolites detected in the cecum of *ApoE* KO mice colonized with communities from mice with disparate atherosclerosis phenotypes**

**Supplementary Table 10. Strains and primers used in this study.**

