## Supplemental Figures for "Gut bacteria impact host uric acid burden and its association with atherosclerosis"

Supplementary Figure 1

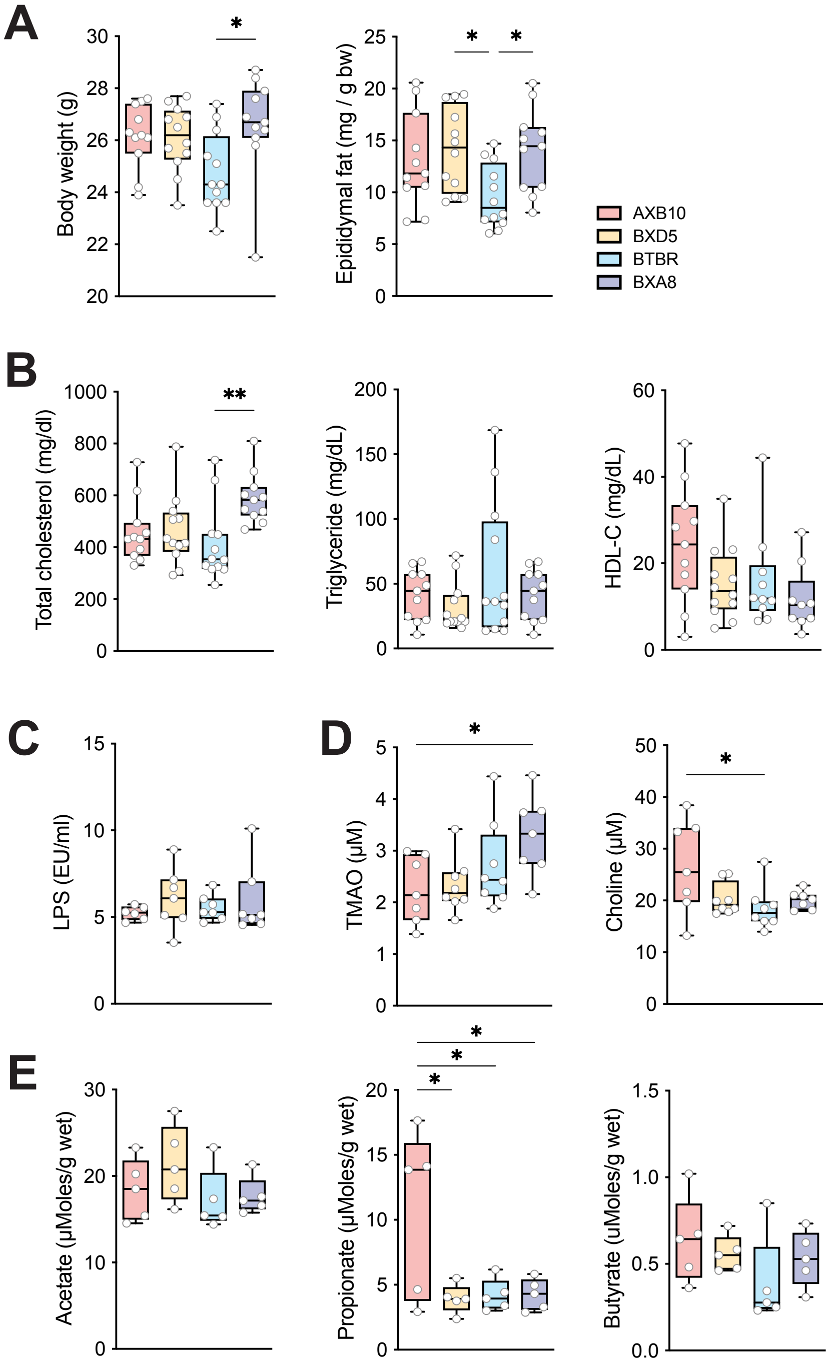

### Supplementary Figure 2

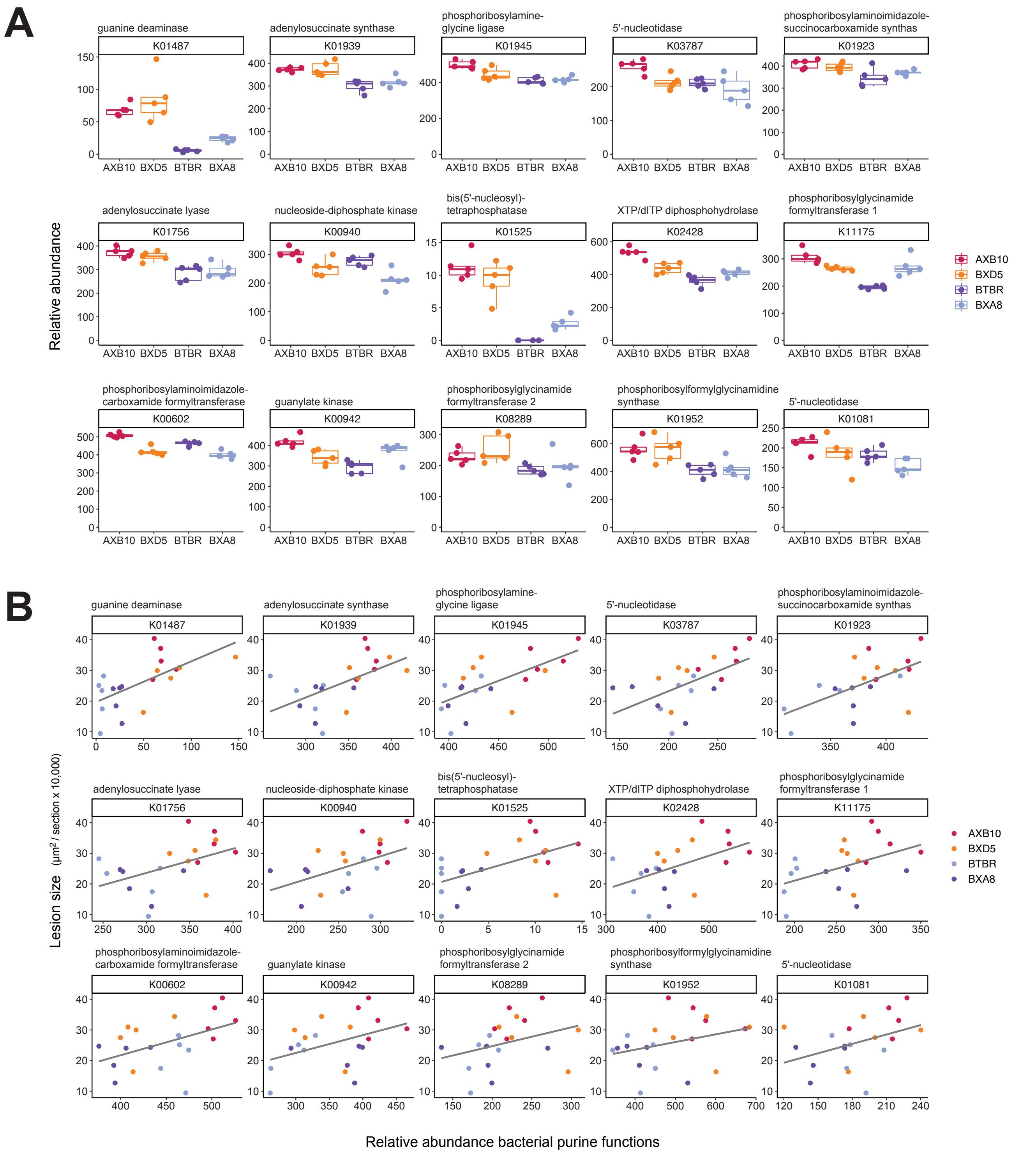

### Supplementary Figure 3

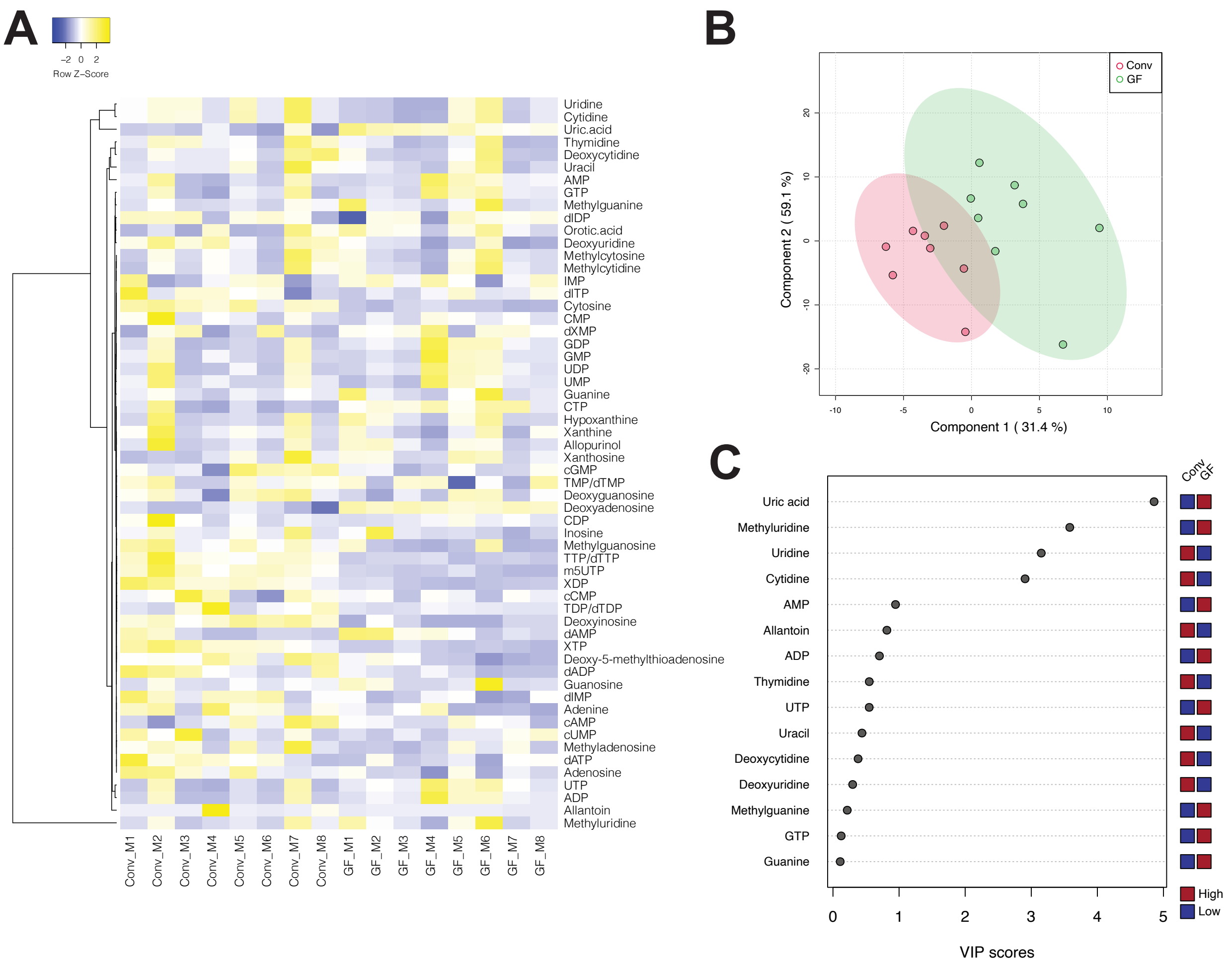

### Supplementary Figure 4

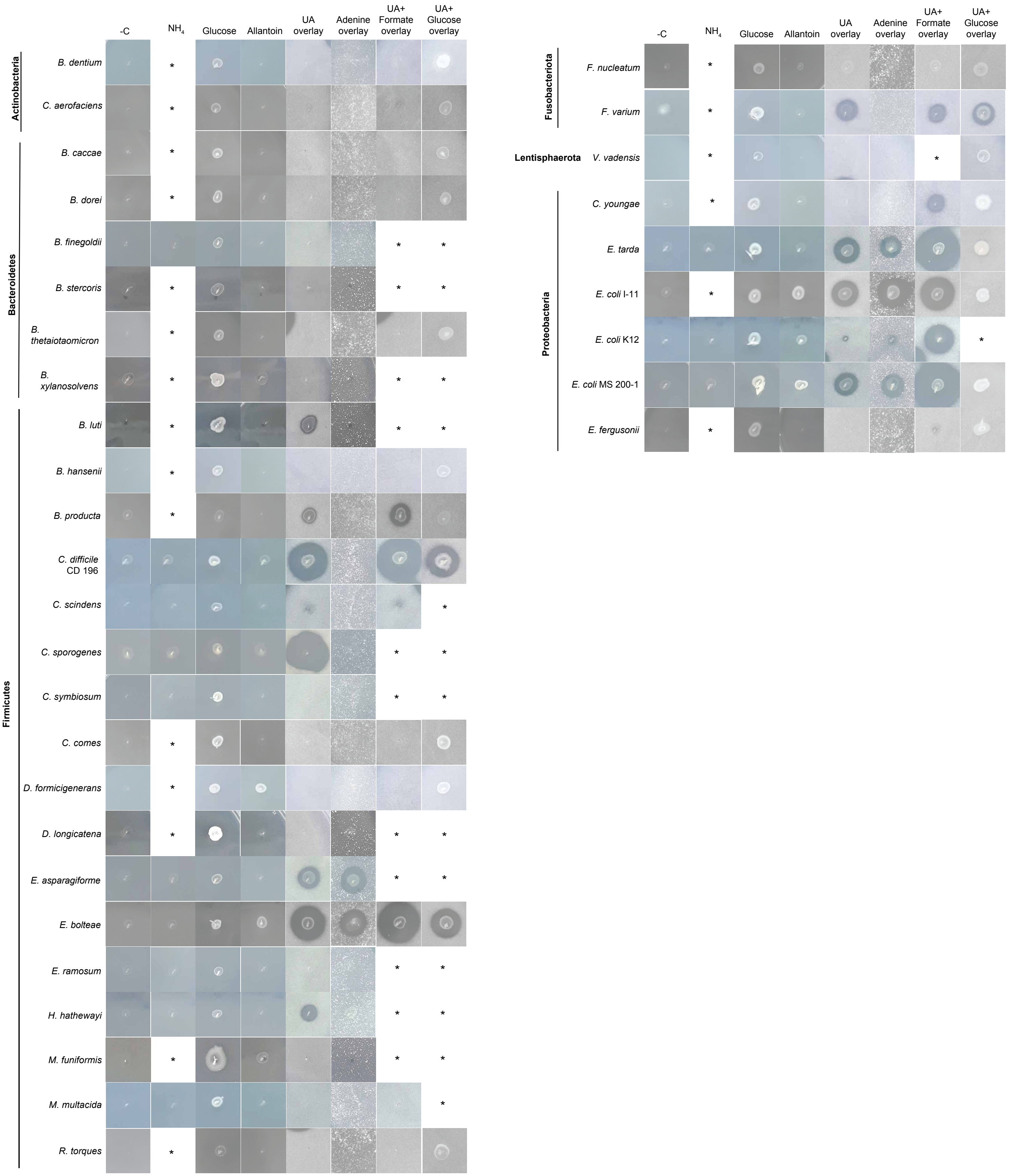

Supplementary Figure 5

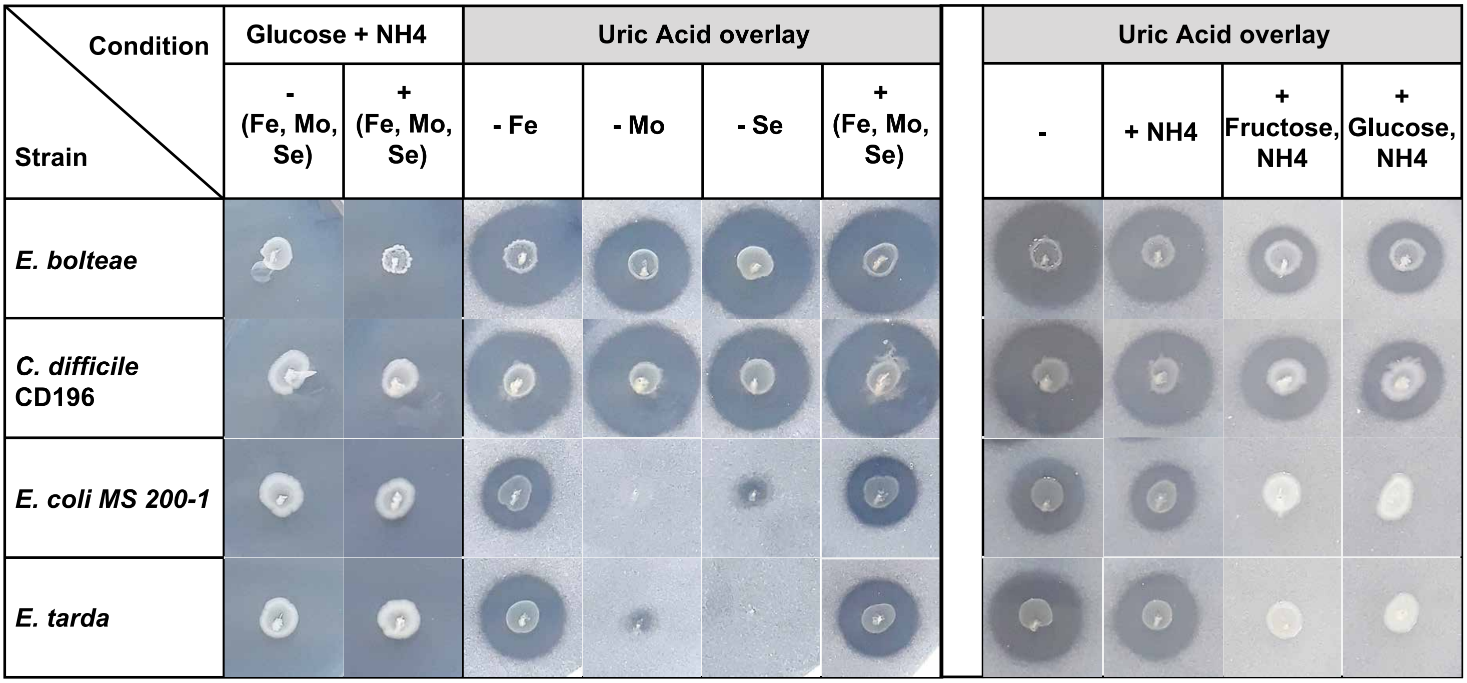

### Supplementary Figure 6

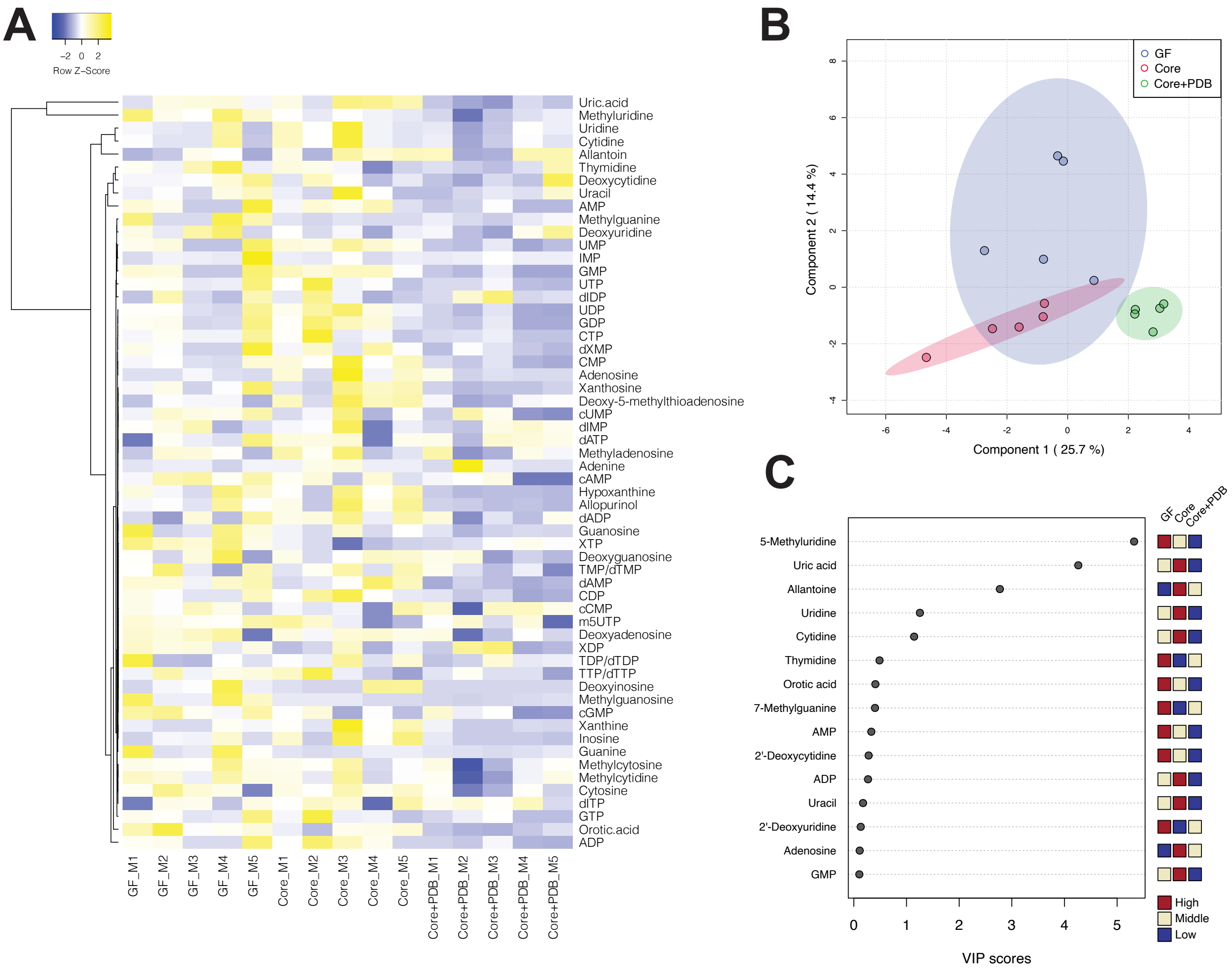

### Supplementary Figure 7

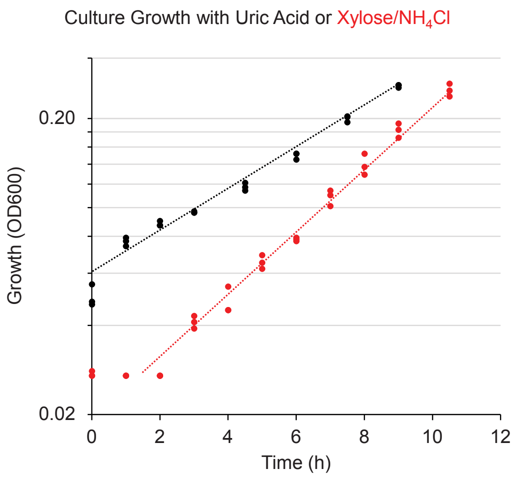

### Supplementary Figure 8

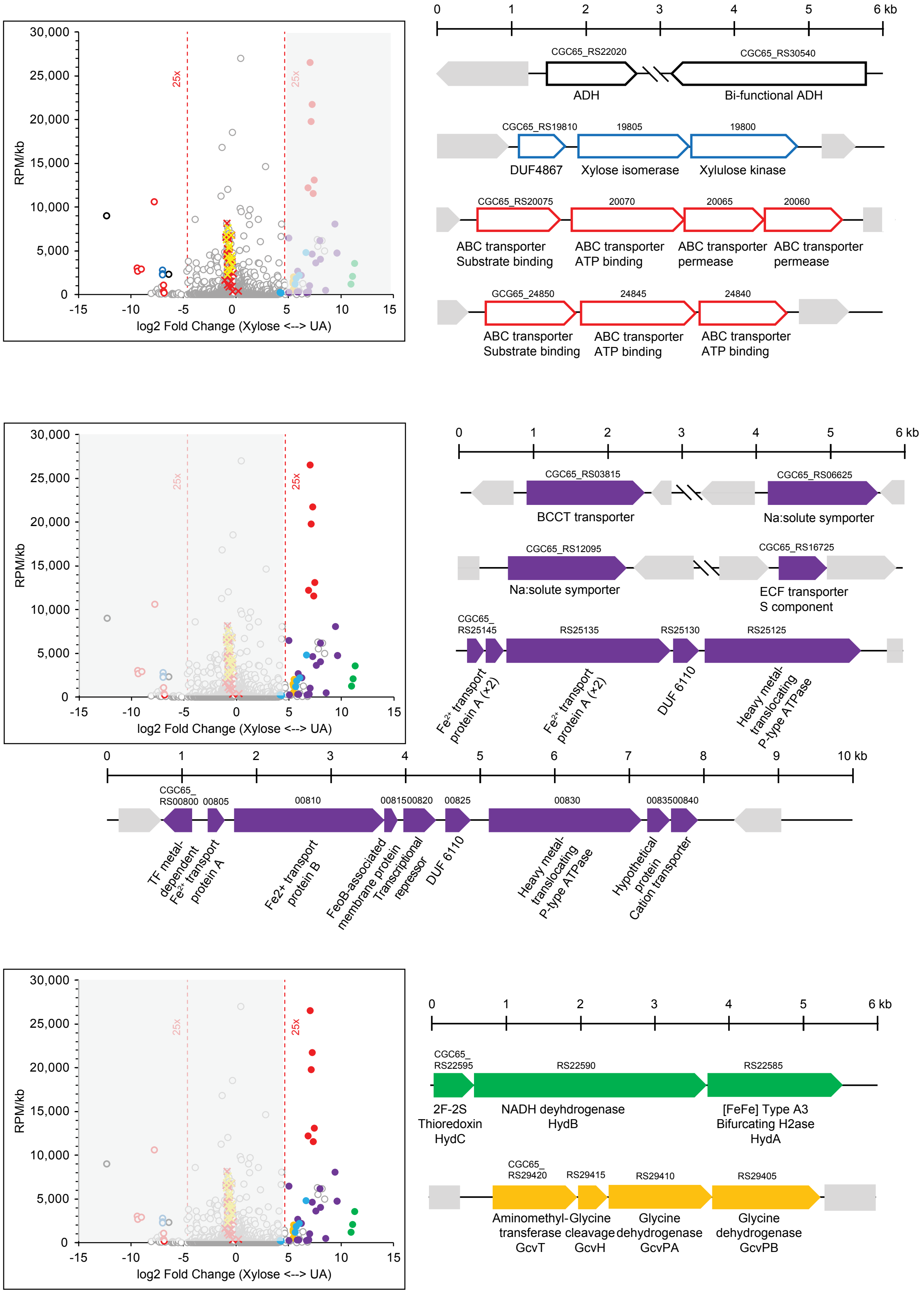

Supplementary Figure 9

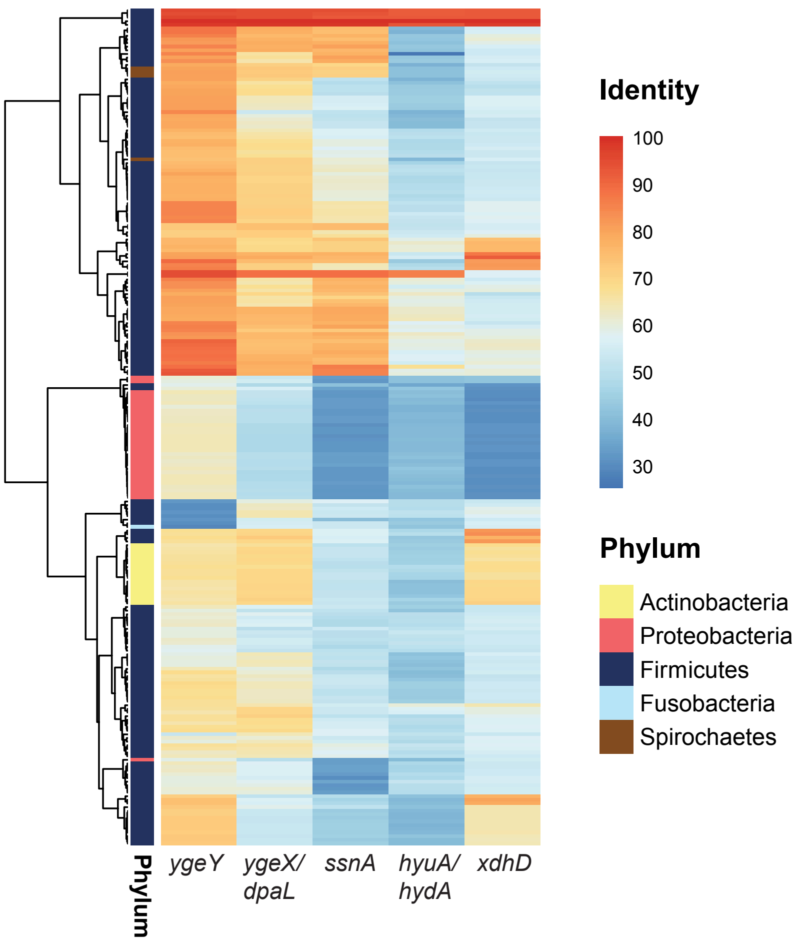
