## Supplemental Methods for "Gut bacteria impact host uric acid burden and its association with atherosclerosis"

**Supplementary Methods**

Media formulations

**METHOD:** **#23B medium (2×) for liquid cultures**

| **Component** |  | **Amount** | **[Medium]** | **Notes/Comments** |
| --- | --- | --- | --- | --- |
| Milli-Q H_2_O  1 M K_x_PO_4_, pH 7.2  0.025% Resazurin  Yeast Extract  NaAcetate·3H_2_O FW 136  NaCl FW 58.44  0.5 M K_2_SO_4_  1.0 M MgCl_2_·6H_2_O  1.0 M CaCl_2_·2H_2_O  NaHCO_3_ FW = 84.0  “2x” Vit. K_1_ + K_3_ soln.  ATCC Vitamin Mixture  (MD – VS)  - - - - - - - - - - - - -  Adjust pH @ 7.0  ~600 ul 10N NaOH 🡪  pH 7.23  - - - - - - - - - - - - -  “50×” Trace Minerals  5 mM FeSO_4_·7H_2_O  Add Milli-Q H_2_O to:  L-cysteine·HCl FW 174.63 | - -  - - | 400 ml  50 ml  2.0 ml  1.0 g  3.4 g  1.46 g  1.0 ml  1.0 ml  0.5 ml  1.68 g  125 µl  10 ml  - - - - - - -  - - - - - - -  5.0 ml  1.0 ml  500 ml  0.88 g | **per 1×**  50 mM  0.10%  25 mM  25 mM  0.5 mM  1 mM  0.5 mM  20 mM  - - - - -  - - - - -  (1:200)  5 µM  5 mM | - - - - - - - - - - - -  - - - - - - - - - - - - |
| Immediately filter sterilize medium; place in anaerobic glove chamber, loosen cap overnight, retighten cap.  To prepare cultures combine 1:1 with anerobic, sterile water. Add stocks of carbohydrates and NH_4_Cl as required.  To prepare cultures for purine catabolism, introduce purine (e.g., 12 mg uric acid) into 125 × 16 mm Hungate tubes, transfer into the anaerobic chamber and allow deoxygenation (overnight). Then add 5 ml Milli-Q purified anaerobic water, seal tubes and autoclave (121°C/20 minutes). Vortex vigorously when removed from the autoclave to disperse substrate, allow to cool, then introduce 5 ml sterile medium/tube by syringe. | | | | |

**METHOD:** **#26B medium for plates**

| **Component** |  | **Amount** | **[Medium]** | **Notes/Comments** |
| --- | --- | --- | --- | --- |
| Milli-Q H_2_O  1 M K_x_H_x_PO_4_, pH 7.2  0.025% Resazurin  Tricine FW 179.2  Yeast Extract  NaAcetate·3H_2_O FW 136  NaCl FW 58.44  0.5 M K_2_SO_4_  1.0 M MgCl_2_·6H_2_O  1.0 M CaCl_2_·2H_2_O  NaHCO_3_ FW = 84.0  “2x” Vit. K_1_ + K_3_ soln.  ATCC Vitamin Mixture  (MD – VS)  - - - - - - - - - - - - -  Adjust pH To 7.3(@6.99)  ~ 600 ul 10N NaOH 🡪pH 7.27  - - - - - - - - - - - - -  Add Milli-Q water to:  “500×” TM #4  5 mM FeSO_4_·7H_2_O  5 mM Mo  1 mM Se  L-cysteine·HCl FW 175.64 | -  - | 800 ml  50 ml  2.0 ml  0.72 g  2.0 g  3.4 g  1.46 g  1.0 ml  1.0 ml  0.5 ml  1.68 g  126 µl  10 ml  - - - - - - -  - - - - - - -  1000 ml  2.0 ml  1.0 ml  2.0 ml  1.0 ml  0.88 g | 50 mM  4 mM  0.20%  25 mM  25 mM  0.5 mM  1 mM  0.5 mM  20 mM  - - - - -  - - - - -  5 µM  10 µM  1 µM  5 mM | [Plates = Medium × ½]  25 mM  2 mM  0.10%  12.5 mM  12.5 mM  0.25 mM  0.50 mM  0.25 mM  10 mM  - - - - - - - - - - - -  - - - - - - - - - - - -  2.5 µM  5.0 µM  0.5 µM  2.5 mM |
| Immediately filter sterilize medium; place in anaerobic glove chamber, loosen cap overnight, retighten cap.  Combine in the ratio of 1:1 with sterile, molten 2.4% Bacto Agar. | | | | |

**METHOD:** **#26B-All medium for monolayer plates containing allantoin**

| **Component** |  | **Amount** | **[Medium]** | **Notes/Comments** |
| --- | --- | --- | --- | --- |
| Milli-Q H_2_O  1 M K_x_H_x_PO_4_, pH 7.2  0.025% Resazurin  Tricine FW 179.1  Yeast Extract  NaAcetate·3H_2_O FW 136  NaCl FW 58.44  0.5 M K_2_SO_4_  1.0 M MgCl_2_·6H_2_O  1.0 M CaCl_2_·2H_2_O  NaHCO_3_ FW = 84.0  “2x” Vit. K_1_ + K_3_ soln.  Allantoin FW 158.1  ATCC Vitamin Mixture  (MD – VS)  - - - - - - - - - - - - -  Adjust pH To 7.3(@7.06)  ~ 260 ul 10N NaOH 🡪 pH 7.27 |  | 440 ml  16.7 ml  0.67 ml  0.24 g  0.67 g  1.14 g  0.49 g  0.333 ml  0.333 ml  0.167 ml  0.56 g  42 ul  4.74 g  3.33 ml  - - - - - - - | 33.4 mM  2.68 mM  0.134%  16.7 mM  16.7 mM  0.333 mM  0.667 mM  0.334 mM  13.33 mM  60 mM  - - - - - | [Plates = Medium × 3/4]  25 mM in plates  2 mM  0.10%  May omit  12.6 mM  0.25 mM  0.50 mM  0.25 mM  10 mM  45 mM (Alfa Aesar A15571.30 lot 10199372, 98%)  - - - - - - - - - - - - |
| Adjust pH to 7.3 (@ 7.06): add ~ 260 µl 10N NaOH, pH --> 7.27  After pH adjustment heat to ~50°C with mixing to fully dissolve Allantoin  (microwave 3×30 seconds). | | | | |
| Add Milli-Q water to:  “500×” TM #4  5 mM FeSO_4_·7H_2_O  5 mM NaMoO_4_·2H_2_O  1 mM Na_2_SeO_3_  L-cysteine·HCl FW 175.64 |  | 500 ml  0.667 ml  0.333 ml  0.667 ml  0.333 ml  0.30 g | 3.33 µM  6.67 µM  0.67 µM  3.42 mM | 2.5 µM  5.0 µM  0.5 µM  2.6 mM |
| Immediately filter sterilize medium; place in anaerobic glove chamber, loosen cap overnight, retighten cap.  Combine in the ratio of 1:3 with sterile, molten 4.8% Bacto Agar.  Note: upon prolonged storage some allantoin will recrystallize. | | | | |

**METHOD:** **11E rich medium with glucose + maltose** **(2X)**

| **Component** |  | **Amount** | **[1× Medium]** | **Notes/Comments** |
| --- | --- | --- | --- | --- |
| tap distilled H_2_O  1 M K_x_H_x_PO_4_, pH 7.2  Tryptone  Yeast Extract  Meat Extract  Tween 80 (25% soln.)  0.025% Resazurin  NaAcetate∙3H_2_O  D-(+)-Glucose  FW = 180.16  Maltose FW = 342.3  [L-Lysine·2HCl  FW = 219.11  __ ______________________  NaHCO_3_ FW = 84.0  Histidine-Hematin soln.  ATCC Vitamin soln.  - - - - - - - - - - - - -  Adjust pH to ~7.2 + ~350 µl 10N NaOH  - - - - - - - - - - - - -  5 mM FeSO_4_·7H_2_O  Adjust volume = **250 ml**  L-cysteine |  | 200 ml  30 ml  8.0 g  4.0 g  4.0 g  0.5 ml  1.0 ml  1.7 g  1.35 g  1.71  2.75 g]  ________  1.05 g  0.25 ml  2.5 ml  - - - - - - -  - - - - - - -  0.25 ml  0.25 g | 60 mM  25 mM  15 mM  10 mM  25 mM  _______  25 mM  - - - - -  - - - - -  2.5 µM | BD 211921  Fisher Sci. BP1422  HiMedia RM003  >Optional  >As required  >Other C source(s) as necessary  - - - - - - - - - - - -  - - - - - - - - - - - - |
| Immediately filter sterilize medium; place in anaerobic glove chamber, loosen cap overnight, retighten cap.  1x medium: Combine with equal volume sterile anaerobic water.  Plates: Combine with equal volume 2.4% agar. | | | | |

Stock solutions:

1M Potassium Phosphate Buffer, pH 7.2

121.9 g K_2_HPO_4_ (anhydrous, MW = 174.2) + 40.8 g KH_2_PO_4_ (anhydrous MW = 136.1) dissolved in ~900 ml Milli-Q H_2_O. Check pH (~7.2), adjust pH to 7.2, adjust volume to 1.0 l.

5 mM FeSO_4_·7H_2_O (Fisher Scientific #I146-500)

0.139 g in 100 ml H_2_SO_4_-acidified MilliQ water (acidify with 4 drops H_2_SO_4_).

5 mM Na_2_MoO_4_·2H_2_O (MP Biomedicals #194863)

0.121 g in 100 ml H_2_SO_4_-acidified MilliQ water.

1 mM Na_2_SeO_3_ (Acros Organics #200730250)

0.0173 g in 100 ml H_2_SO_4_-acidified MilliQ water.

“2×” Vitamins K_1_ + K_3_ solution

0.1 ml viscous liquid vitamin K_1_ (Chem-Impex Intl., #00768) dissolved into 10 ml 100% ethanol, + 20 mg vitamin K_3_ (Sigma M5625), filter sterilize, store at -20°C.

Histidine-Hematin solution

1. 0.2 M histidine, pH 8.0: Add 2.1 g histidine-HCl·H_2_O (Sigma #H7875) into 40 ml distilled H_2_O. Adjust pH from ~4 to 8.0 with 10N NaOH (just over 1 ml, histidine should dissolve). Adjust the final volume to 50 ml with distilled H_2_O.

2. Mix 12 mg hematin (Sigma #H3281) + 10 ml 0.2 M histidine, pH 8.0. Dissolve by vigorous shaking for several hours. Filter sterilize, store aliquots at -20°C.

“50×” Trace Minerals (TM)

1000 ml MilliQ water, acidified with 5 drops H_2_SO_4_. Store under N_2_, refrigerated.

| **Component** | **FW** | **g** | **[Stock]** | **Medium**  **(1:200-diluted)** |
| --- | --- | --- | --- | --- |
| MnCl_2_·4H_2_O | 197.9 | 0.2969 | 1500 µM | 7.5 µM |
| ZnCl_2_ | 136.3 | 0.0682 | 500 | 2.5 |
| CoCl_2_·6H_2_O | 237.9 | 0.0476 | 200 | 1.0 |
| Na_2_MoO4·2H_2_O | 242.0 | 0.0121 | 50 | 0.25 |
| Na_2_SeO_3_ | 172.9 | 0.0086 | 50 | 0.25 |
| NiCl_2_·6H_2_O | 237.7 | 0.0594 | 250 | 1.25 |
| Na_2_WO_4_·2H_2_O | 329.9 | 0.0165 | 50 | 0.25 |

Trace Minerals (TM) #4

1000 ml MilliQ water, acidified with 5 drops H_2_SO_4_. Store under N_2_, refrigerated.

| **Component** | **FW** | **g** | **5000×[Stock]** | **500×[Stock]** | **Medium**  **(1:500-diluted)** |
| --- | --- | --- | --- | --- | --- |
| MnCl_2_·4H_2_O | 197.9 | 0.0990 | 500 µM | 50 µM | 100 nM |
| ZnCl_2_ | 136.3 | 0.0682 | 500 | 50 | 100 |
| CoCl_2_·6H_2_O | 237.9 | 0.0595 | 250 | 25 | 50 |
| NiCl_2_·6H_2_O | 237.7 | 0.1188 | 500 | 50 | 100 |
| Na_2_WO_4_·2H_2_O | 329.9 | 0.0824 | 250 | 25 | 50 |

Preparation of top layer overlays for bilayer plates:

**Uric Acid top layer, ~7 ml/plate**

20 ml tap distilled H_2_O plus:

1.0 ml of 1M stock potassium phosphate buffer, pH 7.2

0.48 g Uric Acid (Sigma-Aldrich U0881-100G Pcode 102408543, ≥99%)

pH measured at 7.0

0.56 g Bacto Agar (in 40 ml, = 1.4%)

Autoclave 20 min./121°C, mix rapidly to evenly disperse uric acid, under anaerobic conditions add 20 ml medium 26B, mix well and transfer 7 ml onto base layer medium (7 ml = 84 mg uric acid). Mix between each addition to keep suspension uniform.

**Adenine top layer, ~7 ml/plate**

20 ml tap distilled H_2_O plus:

1.0 ml of 1M stock potassium phosphate buffer, pH 7.2

0.55 g Adenine (Alfa Aesar A14906, 99%)

pH measured at 7.2

0.56 g Bacto Agar (in 40 ml, = 1.4%)

Autoclave 20 min./121°C, mix rapidly to evenly disperse adenine, under anaerobic conditions add 20 ml medium 26B, mix well and transfer 7 ml onto base layer medium (7 ml = 84 mg uric acid). Mix between each addition to keep suspension uniform.
